# DNA-Diffusion: Leveraging Generative Models for Controlling Chromatin Accessibility and Gene Expression via Synthetic Regulatory Elements

**DOI:** 10.1101/2024.02.01.578352

**Authors:** Lucas Ferreira DaSilva, Simon Senan, Zain Munir Patel, Aniketh Janardhan Reddy, Sameer Gabbita, Zach Nussbaum, César Miguel Valdez Córdova, Aaron Wenteler, Noah Weber, Tin M. Tunjic, Talha Ahmad Khan, Zelun Li, Cameron Smith, Matei Bejan, Lithin Karmel Louis, Paola Cornejo, Will Connell, Emily S. Wong, Wouter Meuleman, Luca Pinello

## Abstract

The challenge of systematically modifying and optimizing regulatory elements for precise gene expression control is central to modern genomics and synthetic biology. Advancements in generative AI have paved the way for designing synthetic sequences with the aim of safely and accurately modulating gene expression. We leverage diffusion models to design context-specific DNA regulatory sequences, which hold significant potential toward enabling novel therapeutic applications requiring precise modulation of gene expression. Our framework uses a cell type-specific diffusion model to generate synthetic 200 bp regulatory elements based on chromatin accessibility across different cell types. We evaluate the generated sequences based on key metrics to ensure they retain properties of endogenous sequences: transcription factor binding site composition, potential for cell type-specific chromatin accessibility, and capacity for sequences generated by DNA diffusion to activate gene expression in different cell contexts using state-of-the-art prediction models. Our results demonstrate the ability to robustly generate DNA sequences with cell type-specific regulatory potential. DNA-Diffusion paves the way for revolutionizing a regulatory modulation approach to mammalian synthetic biology and precision gene therapy.

## Introduction

Gene regulation is a complex process orchestrated at different levels: genetic, epigenetic, and post-transcriptional. The genomic DNA encodes the blueprint for proteins and the regulatory elements that control when, where, and how much of each protein is made. These regulatory elements, such as promoters, enhancers, silencers, and insulators, interact with various proteins and RNA molecules to modulate the transcriptional activity of genes. Unraveling the intricacies of regulatory elements and harnessing their power for gene expression control are defining pursuits in genomics and synthetic biology. However, the systematic modification and optimization of these elements to achieve desired expression patterns remains a major hurdle. This process can potentially correct disease-related misregulation and direct cells to specific functional states.

Despite the availability of technologies to annotate regulatory elements, thanks to the efforts of large consortia such as ENCODE^12^, Roadmap Epigenomics^2,3^, Blueprint ^4^, FANTOM^4,5^ and others, and the ability to understand their critical nucleotides through techniques like MPRA (Massively Parallel Reporter Assays) and CRISPR-based perturbations, there remains a significant challenge in fully comprehending these regulatory elements. These consortia have uncovered the complexity of gene regulation and provide rich data sources for learning about regulatory element features. However, to gain a deeper insight into DNA regulatory grammar, it is crucial to demonstrate how to synthesize these elements *de novo*.

The generative artificial intelligence (AI) field has advanced tremendously over the last few years, yielding approaches that enable researchers to discover, represent, and generate patterns in biological data like never before. These AI-driven methods can be pivotal in understanding the nuances of gene regulation. By using AI to design synthetic sequences and identify genomic locations for their integration, researchers can experiment with and observe the effects of these synthetic elements in natural biological systems. These tools can help confirm the roles of various regulatory elements and safely and precisely modulate gene expression. The synthesis of regulatory elements *de novo*, guided by generative AI models, represents a promising frontier in understanding and manipulating the complex gene regulatory networks underlying and cell states.

Diffusion models have emerged as a powerful class of generative models that have achieved strong performance in generating synthetic images, audio, and text^6–9^ and more recently also adapted in the biological context to generate protein structures^10^ (**Fig. 1a**). This adaptability in creating diverse outputs across different fields has inspired their exploration in additional scientific areas. In this manuscript, we propose investigating the generation of synthetic DNA regulatory elements through these models, assessing their applicability and efficacy in the specific context of DNA sequence generation.

**Figure 1.**
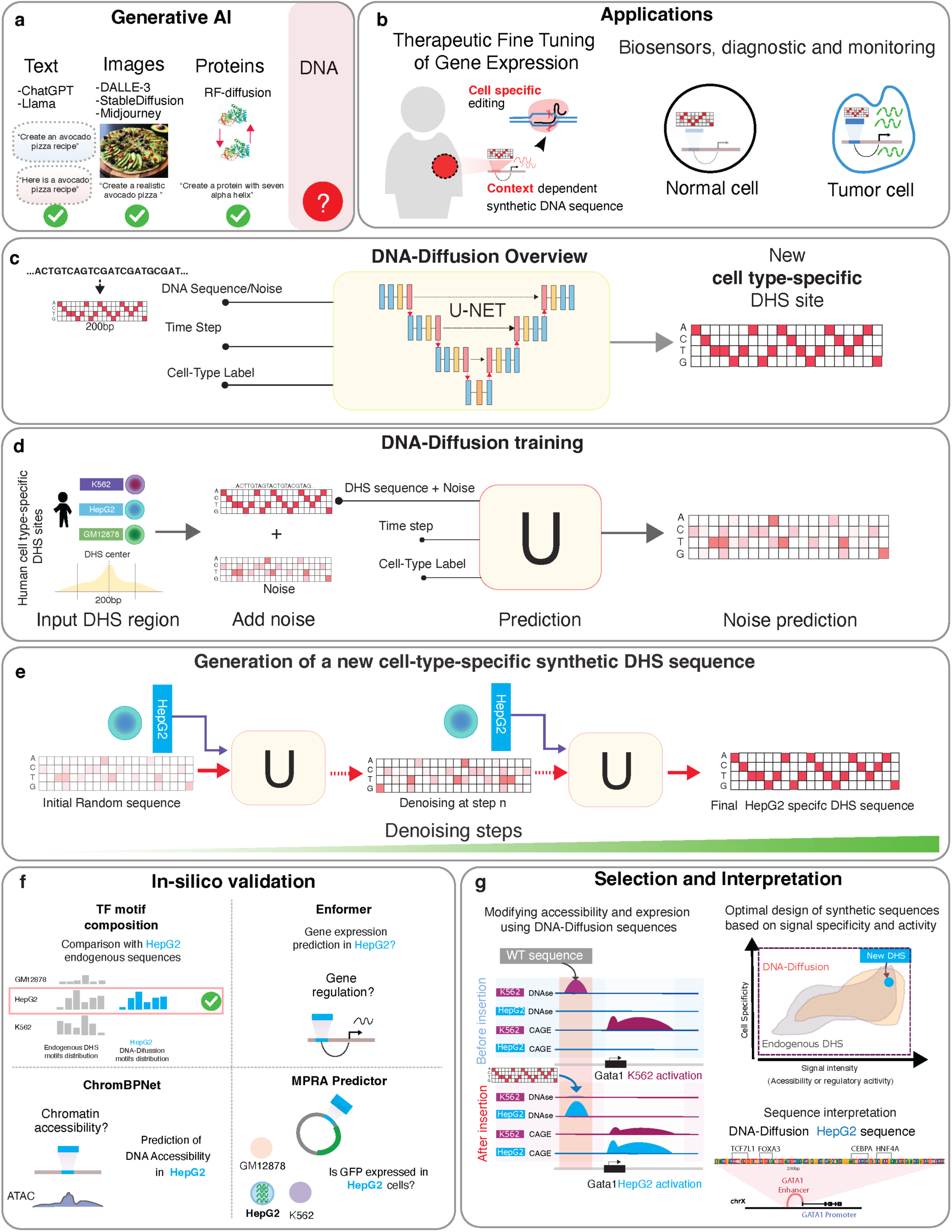
a) Examples of generative AI models applied to language, image generation, and biological problems. b) Exploration of potential therapeutic applications based on synthetic cell-type specific sequences. c) Schematic overview of the DNA-Diffusion model. The model utilizes a U-Net architecture to generate new DNA sequences iteratively based on cell types presented in the training dataset. d) The DNA-Diffusion model was trained using unique DHS DNA sequences from different cell types, including K562, HepG2, and GM12878. The training involves transforming the endogenous DHS sequence into a hot-encoded format and introducing a fixed amount of standard normal noise. The trained U-Net uses the expected noise level (determined by the time step) and cell type information to predict and remove the added noise. This noise prediction process is repeated during training across the entire sequence dataset with varied noise intensities. Once trained, the U-Net can predict the initial noise added to the original DHS endogenous sequences, enabling the generation of new sequences specific to different cell types. e) To generate a new sequence given a cell-type label, a hot-encoded DNA matrix with random Gaussian noise is generated, and the U-Net iteratively denoises this matrix over 50 steps, progressively converging to a sequence that reflects the characteristics of the target cell type. f) Different in silico validations were utilized to evaluate the accessibility, regulatory activity, and motif composition of DNA-Diffusion and endogenous DHS regions. g) Framework developed for selecting and interpreting generated sequences based on cell-type signal specificity, intensity, or motif composition.

Diffusion models operate by taking an input pattern in a specific domain, such as an image, and progressively adding Gaussian noise until the pattern is indistinguishable from noise (forward process). The training of these models involves inverting this process, learning to remove the noise across various timesteps to recreate the original pattern (reverse process). A denoising diffusion probabilistic model^7^ (DDPM) is one formulation for diffusion models that relies on Markov chains for these two processes and provides the foundation for our research. This reverse process, capable of reconstructing patterns from noise, enables the model to generate new samples that mirror the data distribution of the training set, effectively creating novel patterns from pure noise.

Recent advances in synthetic DNA sequence design have utilized machine learning models such as Generative Adversarial Networks (GANs) and Convolutional Neural Networks (CNNs), which have demonstrated potential in sequence generation ^11–13^. Initial efforts have been starting to explore diffusion models^14,15^ without employing conditional diffusion, therefore limiting the generation of cell type-specific regulatory elements. All these methods typically require an orchestrated effort involving training multiple models and complex layers, including external predictors or classifiers, to tailor the design process to specific cell types or motif compositions. This orchestration underscores a significant limitation: the inability to train these interconnected models simultaneously and end-to-end for multiple cell types. The complexity arises not only from the need for multiple models, but also from the intricate layers and heuristics necessary to synchronize and fine-tune predictions for each distinct cell type. A unified approach is essential to expand and effectively train these models across hundreds of cell types, leveraging the wealth of available data. Such an approach could pave the way for foundational models in DNA sequence design, capable of accommodating diverse cellular contexts.

Addressing the limitations of current synthetic DNA sequence design methods and exploring the opportunities for the use of diffusion-based approaches, we introduce DNA-Diffusion, a novel approach leveraging diffusion probabilistic models for the design of context-specific DNA regulatory sequences. Unlike other methods, DNA-Diffusion allows training a single model across different cell types and generating cell type-specific sequences without the need for additional models as guidance.

We demonstrate that our model can generate regulatory elements that, in silico, recapitulate endogenous properties like transcription factor (TF) binding motif composition and the ability to reactivate gene expression in a cell-type specific fashion. Further, we discuss strategies to bias generation towards sequences with high signal or high specificity in a given cell type, as well as methods for optimally selecting genomic target regions that can be replaced with these sequences to achieve maximal activity.

Our DNA-Diffusion model bridges the gap between AI-driven generative models and practical applications of synthetic DNA sequence generation for cell type-specific gene regulation. These sequences hold the potential to modify gene expression and be employed in new therapeutic applications requiring precise perturbation of gene regulation (**Fig. 1b**).

## Results

### DNA-Diffusion: a conditional diffusion model to generate cell type-specific regulatory elements

DNA-Diffusion is a conditional diffusion model^16^ that operates in the space of DNA sequences. Sequences are encoded using a strategy akin to one-hot encoding, but each nucleotide has a support range of [-1,1] to facilitate the injection of Gaussian noise centered around zero. The backbone of the model is a denoising U-Net^17^ with two embedding layers for cell label and timestep, respectively (**Fig. 1c)**. During training, it receives three inputs: DNA sequences, a timestep, and cell type labels. After training, the model takes in input a cell type label and can generate novel cell type-specific sequences.

The primary goal in training this model was to generate cell type-specific regulatory elements. We utilized the DHS index dataset curated by Meuleman et al.^18^, which includes 733 biosamples from 438 cell and tissue types, to derive cell type-specific sequences for GM12878, K562, and HepG2. These cell types were chosen for their distinct biological contexts, diverse tissue origins and encompassing different germ layer lineages: GM12878 (a B lymphocyte cell line) for the immune system, K562 (a leukemia cell line) for blood cancer research, and HepG2 (a hepatocellular carcinoma cell line) for liver biology and disease studies. This diverse selection allows for a comprehensive evaluation of the model’s capacity to generate regulatory elements across a wide range of cellular functions and conditions. Additionally, a rich array of genomic tracks, including histone marks, transcription factor binding sites, and expression data, are available for these cell types, providing valuable data for model interpretability and evaluation. Furthermore, the ease of culturing these cell lines in the lab opens avenues for potential future experimental validation of the model’s predictions.

For each cell type, we assigned a categorical class label and selected sequences spanning 200 bp around the summit of cell type-specific peaks (see **Methods**). Prior to model training, we adopted a strategy proposed by Meuleman et al.^19^. Briefly, this chromosome-based stratified sampling strategy was employed to partition the dataset into mutually exclusive subsets for training, validation, and testing. Chromosome 1 (chr1) was designated as the independent test set, chromosome (chr2) was allocated for validation purposes, and the rest of the chromosomes were used for training, which also ensured model generalizability. This enabled conditional training of the DNA-Diffusion model and the generation of cell type-specific sequences. The training involved a forward process where varying levels of noise were introduced into the encoded sequence representations, aiming to learn a function that can effectively denoise a sequence at each step by predicting the patterns of introduced noise (**Fig. 1d**). Upon completing training, a reverse process synthesizes novel DNA sequences, given a specified number of steps and the desired cell type-specific label (**Fig. 1e**).

Having successfully trained the model (**Supplementary** Fig 1), we generated 100,000 DNA sequences per cell type of 200 bp in length. Our goal was to assess the performance of this model architecture across various genomic characteristics: sequence uniqueness, transcription factor composition, chromatin accessibility, and the capacity to drive the expression of nearby genes in a cell type-specific manner (**Fig. 1f,g**), as detailed in the subsequent sections.

### Diffusion-generated sequences exhibit motifs present on endogenous sequences without memorizing existing sequences

To verify the uniqueness of the DNA-Diffusion sequences and confirm that they were not mere copies of the training sequences, we conducted a comparative sequence alignment between the generated DNA-Diffusion sequences and the sets of train, test and randomly sampled endogenous sequences, employing BLAT^20^ analysis for this purpose.

The average BLAT alignment length distribution between the DNA-Diffusion sequences and endogenous training sequences was found to be significantly shorter (23.4bp, t-test p-value < 0.001) than between endogenous test and training sequences (48.8bp) or between randomly sampled genomic sequences and endogenous training sequences (49.3bp) (**Fig. 2c)**. To assess sequence diversity of the generated sequences, we also perform comparative sequence alignment of each set with itself. Comparing all the possible pairwise DNA-Diffusion sequences we observed a significantly lower average match (23bp, t-test p-value < 0.001) as compared to train (29bp) or random (39bp) endogenous sequences **(Fig. 2d)**.

**Figure 2.**
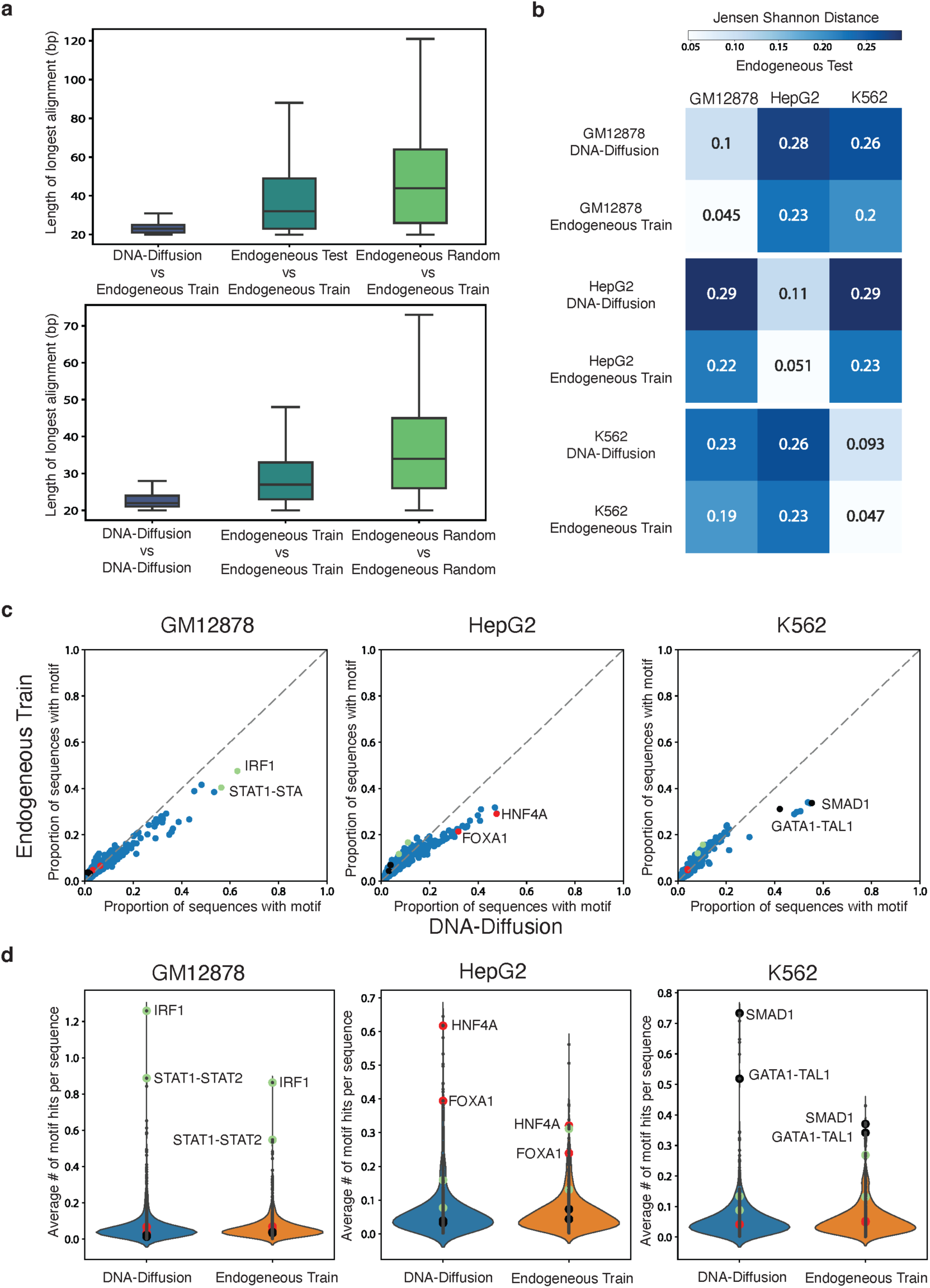
a) Top, bar plots displaying sequence alignment length comparing the amount of overlap between DNA-Diffusion, endogenous test, and endogenous random sequences against endogenous train sequences. Bottom, distribution of alignment length within each group. b) Jensen Shannon distance comparing TF motif hit distributions between DNA-Diffusion sequences and endogenous train sequences against the endogenous test sets across GM12878, HepG2, and K562 cell-types. c) Scatterplot showing the proportion of sequences with a given TF motif in endogenous training sequences vs DNA-Diffusion sequences across GM12878, HepG2, and K562 cell-types. Important cell type-specific TFs are annotated based on existing literature (green: GM12878, red: HepG2, black: K562). d) Violin plot showing the average number of TF motif hits per sequence in endogenous training sequences vs DNA-diffusion sequences across GM12878, HepG2, and K562 cell-types. Important cell type-specific TFs are annotated based on existing literature (green: GM12878, red: HepG2, black: K562).

Among the analyzed DNA-Diffusion sequences, only 111 out of 300k (0.037%) displayed greater than 40bp overlap with the DHS training dataset (52 for GM12878, 30 for HepG2, and 29 for K562). Additionally, assessing the diversity of DNA-Diffusion sequences, we found that of the 300k sequences, only 223 matched with others, with the matches per cell-type being 103 in GM12878 (0.104% of 100,000), 78 in HepG2 (0.079%), and 42 in K562 (0.043%). These findings support that DNA-Diffusion sequences are not copying large sequence segments from the endogenous training set, while also presenting a diverse set of novel sequences.

We next wanted to demonstrate that the diffusion model learned cell type-specific TF composition. Using MOODS ^21^, we scanned the sequences for recognizable TF binding sites using TF position frequency matrices (PFMs) from the JASPAR database^22^. To assess the fidelity of our model’s output to biological reality, we employed the Jensen-Shannon (JS) divergence for comparing the distribution of TF motif hits in cell type-specific DNA-Diffusion sequences against the endogenous test set. This analysis yielded a mean JS divergence of 0.101 across the three cell-types (**Fig. 2c**), revealing a high degree of similarity between the DNA-Diffusion sequences and the endogenous test set motif distributions. Further reinforcing this finding, we established a baseline by comparing endogenous training sequences to endogenous test sequences within each cell type, which showed a closer similarity with a mean JS divergence of 0.048 across the three cell-types (**Fig. 2b**). Importantly, we observed a clear separation of motif composition across cell types in both comparisons, with a slightly more distinct separation in generated sequences (GM12878 DNA-Diffusion: 0.27, GM12878 endogenous train: 0.22, HepG2 DNA-Diffusion: 0.29, HepG2 endogenous train: 0.23, K562 DNA-Diffusion: 0.25, K562 endogenous train: 0.21), suggesting some divergence of TF motif usage from the endogenous sequences.

To elucidate the underlying factors contributing to the observed discrepancy in motif composition between DNA-Diffusion sequences and endogenous training sequences, we compared individual motif abundances across these two sets (**Fig. 2c**). Our analysis revealed that cell type-specific TFs are present in a higher number of DNA-Diffusion sequences compared to endogenous train sequences. For example, HNF4A—a liver-specific pioneer factor ^23^—was detected in approximately 29% of endogenous training sequences, yet it was present in ∼48% of DNA-Diffusion sequences. Similar trends were observed with other liver-specific factors, such as FOXA1 and FOXA3^23,24^. Furthermore, this pattern of enriched cell type-specific TFs in DNA-Diffusion sequences extended to other cell-types. For instance, in the GM12878 cell-type, the immune-related factor IRF1 showed a presence in 63% of DNA-Diffusion sequences, contrasting with 47% in endogenous training sequences. Similarly, in the K562 cell line, the combined GATA1-TAL1 motif ^24,25^ was present in 42% of DNA-Diffusion sequences, compared to 31% in endogenous training sequences.

Interestingly, the DNA-Diffusion sequences exhibited a lower abundance of cell type-specific motifs for cell types other than the targeted one. For instance, IRF1 was found less frequently in HepG2 and K562 specific sequences as compared to the sequences generated for GM12878. This suggests that the model is effectively using cell type-specific motifs to create sequences with greater specificity. On the other hand, endogenous sequences may exhibit a “leaky” or non-specific expression when placed in an alternate cellular context. For example, endogenous sequences selected for HepG2, which more frequently contain the IRF1 motif, might still demonstrate some degree of weak expression when introduced into GM12878 cells, albeit in a less targeted manner.

In addition to the changes in motif abundance across different sets of sequences, we also noted an increase in the average number of motif hits within individual DNA-Diffusion sequences, particularly for cell type-specific transcription factors (**Fig. 2d**). This indicates not just a shift in the overall presence of motifs in the set of DNA-Diffusion sequences, but also an enhanced density of binding sites per sequence. For instance, DNA-Diffusion sequences for HepG2 contained on average about 0.62 HNF4A motifs, in contrast to approximately 0.32 in the endogenous training sequences. In a similar vein, K562 sequences demonstrated an increased number of GATA1-TAL1 and SMAD1 motifs. Likewise, GM12878 sequences were characterized by more frequent occurrences of IRF1 and STAT1 motifs. Overall, this underscores a pattern of increased motif representation per sequence in the DNA-Diffusion sequences across diverse cell types.

In conclusion, these analyses confirm that DNA-Diffusion sequences are not mere replicas of the training set but are instead novel and original sequences. The model enhances cell type-specificity by modulating motif density and incorporating known cell type-specific transcription factors, thereby closely mirroring the motif vocabulary of endogenous sequences for a given cell type. This highlights the model’s capability to generate realistic and diverse genomic sequences for precise cellular contexts without resorting to memorization.

### DNA-Diffusion generated sequences demonstrate cell specificity, accessibility and can activate *cis* genes in a cell type-specific manner *in silico*

To investigate the effects of cell type-specific DNA-Diffusion sequences on chromatin accessibility and gene expression, we replaced endogenous sequences at accessible DHS sites specific to each cell type with our generated DNA-Diffusion sequences. The objective was to determine if these sequences could maintain or change the existing accessibility in their respective cell types. Concurrently, we investigated whether the DNA-Diffusion sequences could induce accessibility in areas that were previously inaccessible, depending on the sequence’s initial cellular context. Additionally, we aimed to evaluate the influence of these sequences on the expression of genes potentially regulated by these elements. To this end we considered different “oracles”, state of the art prediction methods that have demonstrated the ability to recapitulate gene expression and chromatin accessibility patterns across cell types in human cells (**Fig. 3a**).

**Figure 3.**
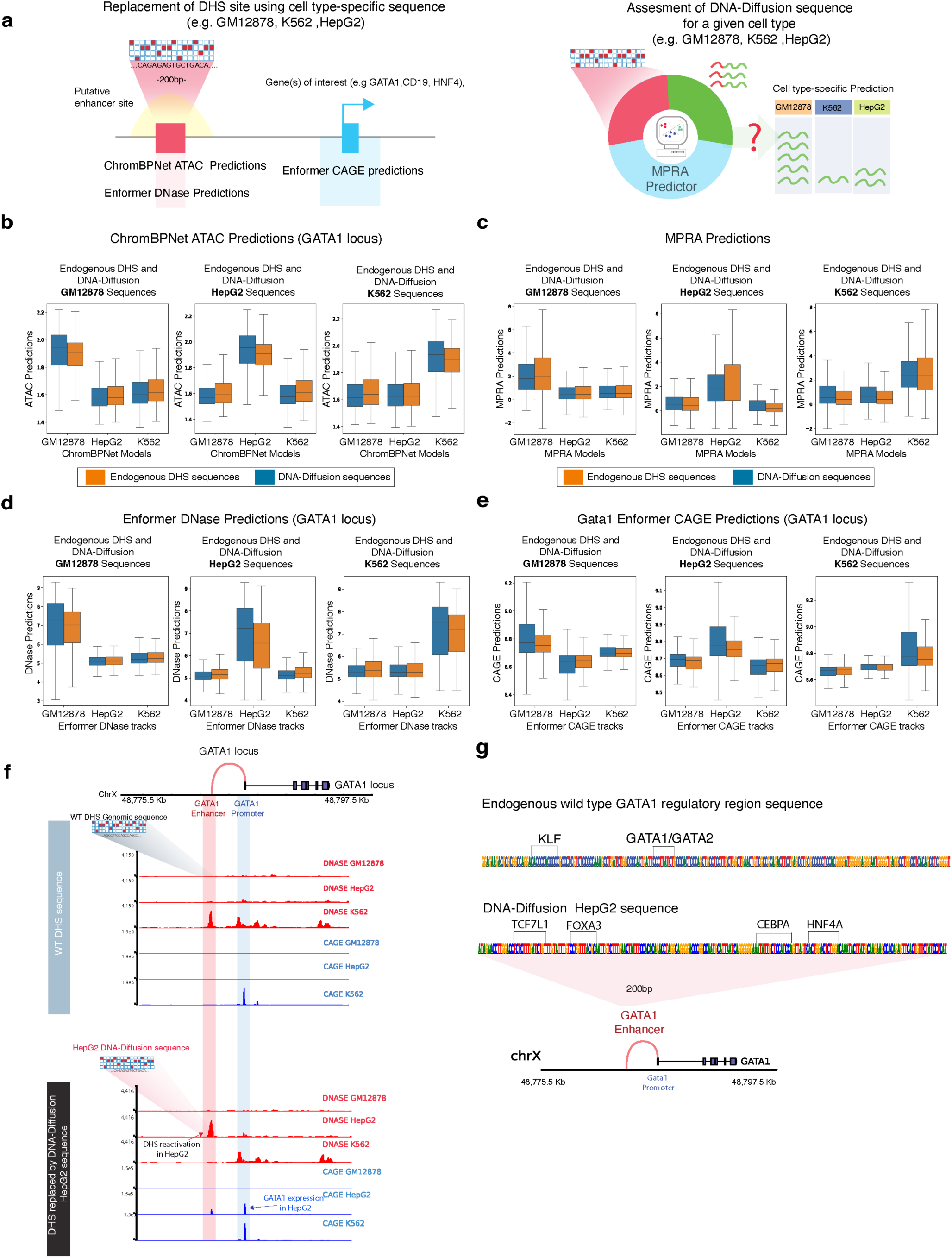
a) Left: Strategy for replacing cell type-specific sequences within putative enhancer regions of target genes (GATA1, CD19, HNF4A), along with predictive modeling of expected chromatin accessibility (ChromBPNet ATAC and Enformer DNase) and regulatory potential (Enformer CAGE). Right: An approach using an MPRA predictor to evaluate the cell type-specific expression potential of the sequences. b) Application of ChromBPNet to predict ATAC accessibility of cell type-specific DNA-Diffusion-generated and endogenous sequences in the GATA1 locus across GM12878, HepG2, and K562 cell lines. c) Measurement of MPRA activity reflecting cell type specificity in DNA-Diffusion and endogenous train sequences for GM12878, HepG2, and K562. d) Enformer DNase predictions of chromatin accessibility at the GATA1 locus, comparing cell type-specific DNA-Diffusion and endogenous train sequences across GM12878, HepG2, and K562. e) Expression of the GATA1 gene driven by both cell type-specific DNA-Diffusion-generated and endogenous train sequences in GM12878, HepG2, and K562. f) Top: DNase and CAGE activity predictions at the GATA1 locus based on the wild-type sequence. Bottom: Post-insertion of a HepG2-optimized DNA-Diffusion sequence into the enhancer region, resulting in heightened chromatin accessibility and GATA1 expression in HepG2, coupled with reduced accessibility in K562; GM12878 shows no substantial alterations. g) Comparative analysis of transcription factor motifs in the wild-type and HepG2-specific DNA-Diffusion sequences of the GATA1 regulatory region, with annotations of established cell type-specific motifs from existing literature.

This approach was applied to the DHS regulatory regions of three distinct and well-known cell type-specific genes for a comprehensive analysis (**Supplementary Table 1**). Specifically, this included GATA1, a transcription factor that plays an essential role in red blood development and is implicated in various blood-related disorders^26,27^. Importantly, the activity of its proximal enhancers has been validated through CRISPR interference (CRISPRi) in K562 cells ^28,29^. Based on these findings, we selected the region of 200bp that showed the strongest activity. HNF4A, another selected gene, plays a crucial role in triggering the transcriptional response in liver cells and is expressed in HepG2^18,30^. Finally, CD19 is an important gene in B-cell development and serves as a key cellular marker for chimeric antigen receptor (CAR) T-cell therapies ^31^. Lacking experimentally validated enhancers for HNF4A and CD19, we leveraged endogenous chromatin accessibility patterns to identify promising candidate regions for evaluating the impact of our engineering sequences. To delineate a panel of candidate enhancers for each gene, we selected a 200bp region within 100KB of its transcription start site (TSS) and characterized by a pronounced cell type-specific pattern of chromatin accessibility (**Supplementary** Fig 2).

Having selected the genomic location of these distal regulatory sequence for each locus in different cellular contexts, we next assessed the impact of replacing the endogenous sequences with different cell type-specific sequences for each cell type from two groups: either from DNA-Diffusion sequences or endogenous DHS from other locations. Our objective was to evaluate the extent to which DNA-Diffusion sequences or endogenous sequences, presumed to be active in a particular cellular context, exhibit cell type-specific patterns upon integration into the regulatory regions of these loci. To this end, we utilized ChromBPNet^32^, a recent state-of-the-art method capable of predicting chromatin accessibility (ATAC signal) at nucleotide level resolution based exclusively on DNA sequence information. We concurrently utilized three distinct ChromBPNet models, each specifically trained on K562, GM12878, and HepG2 data to analyze the predicted chromatin accessibility patterns of each sequence in the various groups **(Methods)**.

Initially, we focused on the GATA1 locus, replacing the endogenous sequence of the validated enhancer (**Sup Fig. 2a**), known for its cell type-specific chromatin accessibility in K562, with sequences from our two groups. Our observations using the three ChromBPNet models as oracles indicated that only DNA-Diffusion and endogenous sequences specific to K562 successfully maintained or enhanced accessibility in K562. As a baseline, predictions from endogenous DHS sites in the training set and specific for K562 cells had a mean log-normalized predicted ATAC value of 1.88 based on the K562 model, compared to 1.65 for the HepG2 model and 1.66 for the GM12878 model. Notably, the predictions for the DNA-Diffusion sequences showed a value of 1.91 for the K562 model, suggesting slightly higher but significant activity (p < 0.01, one-sided t-test) when compared to the endogenous baseline values and slightly lower but significant activity (p < 0.01, one-sided t-test) for the other two cells type models, both showing a predicted value of 1.64 (**Fig 3b**). These results indicate that synthetic K562-specific sequences demonstrated the ability to maintain or enhance chromatin accessibility in K562 cells compared to the endogenous sequence. Conversely, these sequences exhibit reduced accessibility in non-target cell types, suggesting increased context-specificity. Contrasting with the K562 cells, where an established DHS signature demarcates the chosen GATA1 regulatory region **(Fig. 3b)**, HepG2 and GM12878 cells typically display a lack of accessibility at this element (**Supplementary** Fig. 2 **b,c**).

This distinction provided an opportunity to evaluate whether DNA-Diffusion sequences, specifically crafted for these cell types, could effectively reactivate chromatin accessibility in regions usually inactive. Despite the inherent inactivity of the selected region in these cell types, both DNA-Diffusion and endogenous sequences specific to GM12878 and HepG2 managed to successfully induce chromatin accessibility (Fig 3b). Similar to the observations in K562 cells, the DNA-Diffusion sequences for HepG2 and GM12878 showed stronger signal in their respective ChromBPNet models compared to predictions in other cell types. Specifically, DNA-Diffusion sequences for the GATA1 regulatory region in HepG2 had a predicted mean ATAC signal of 1.92, with lower values in GM12878 and K562 models (1.60 and 1.63, respectively). Conversely, GM12878 DNA-Diffusion sequences in the GATA1 regulatory region exhibited a mean predicted value of 1.91 in the GM12878 ChromBPNet model, compared to 1.60 in HepG2 and 1.63 in K562 (**Fig 3b**).

Collectively, these results underscore a context-dependent functionality of DNA-Diffusion sequences, with an enhanced activity within the appropriate cell type and diminished activity in others. Analysis using cell type-specific signals consistently showed that DNA-Diffusion sequences designed for K562, HepG2, and GM12878 exhibited higher predicted accessibility values within their respective models, as opposed to predictions from models trained on different cell types. This evidence supports the cell type-specific utility of the DNA-Diffusion strategy, demonstrating its tailored activation capacity in designated cellular environments.

Extending our investigation to assess the generalizability of our approach, we explored its applicability to the regions selected for the HNF4A and CD19 loci, with the former being accessible in HepG2 but not in K562 and GM12878, and the latter accessible in GM12878 but not in K562 and HepG2 (**Supplementary** Fig 2b,c). Using the same approach, we observed that DNA-Diffusion sequences were able to maintain, augment, or initiate chromatin accessibility across various cell types at these loci, which illustrates these effects in different genomic contexts (**Supplementary** Figs. 3a, 4a). These observations suggest that DNA-Diffusion sequences can achieve a spectrum of chromatin accessibility that is comparable or greater than that of endogenous sequences in the right cellular type and in different genomic contexts. The DNA-Diffusion model not only preserved and modified accessibility in regions with established DHS activity specific to a cell type but also successfully induced accessibility in genomic regions previously inactive in the respective cell types.

Having established that our synthetic DNA-Diffusion sequences can modulate chromatin accessibility, we aimed to investigate their impact on gene expression, a task that our DNA-Diffusion model was not specifically trained on. Massively Parallel Reporter Assays (MPRA) have been used to measure the cell type-specific ability of a given sequence to drive a reporter gene’s expression in a given cell type and in a genomic context-independent manner. In addition, prior work has shown that the MPRA activity of a sequence can be accurately predicted across a variety of human cell lines^33^ by fine-tuning the Enformer^34^ model. We utilized a similar approach and fine-tuned Enformer to predict MPRA activity in 5 cell lines, including K562, GM12878, and HepG2 cells (**Methods**). We aimed to validate, *in-silico*, cell type-specific expression mediated by our synthetic sequences. Our analysis revealed that sequences designed for a particular target cell showed higher predicted MPRA activity within the corresponding cell compared to predictions for other cell types (**Fig. 3c**). For example, K562 DNA-Diffusion sequence predictions exhibited a mean normalized value of 2.60 according to the K562 MPRA model, markedly higher than the values of 0.68 and 0.67 for the GM12878 and HepG2 models respectively (**Fig.3c**). As a baseline, the predicted activity distribution for endogenous train K562 sequences (2.53) closely aligned with the DNA-Diffusion predictions. Similar patterns emerged when analyzing DNA-Diffusion sequences for HepG2 and GM12878, with sequences specifically tailored for each cell type demonstrating higher predicted MPRA values within their respective models (**Fig.3c)**. While the DNA-Diffusion model was not primarily trained to optimize gene expression regulation, it successfully learned to capture inherent regulatory patterns from the endogenous sequences, thereby enabling cell type-specific modulation of predicted MPRA expression.

Building on the evidence that DNA-Diffusion sequences can modulate in silico chromatin accessibility and drive MPRA expression in a cell type-specific manner as demonstrated by ChromBPNet and MPRA predictive models, we sought to delve deeper into their potential to regulate the expression of specific genes. We focused on the replacement of synthetic sequences at the GATA1, HNF4A, and CD19 loci and assessed the impact on gene expression by modulating chromatin accessibility within the cellular contexts of K562, HepG2, and GM12878. A key goal was to ascertain whether these sequences could activate genes not natively expressed in certain cell types.

To this end, we used Enformer, a cutting-edge deep-learning model that has demonstrated the ability to recapitulate gene expression, chromatin histone modification, and accessibility patterns across cell types using only DNA sequence input. Notably, this model accommodates long-range interactions (up to 100 kb) and can potentially capture the intricate interplay among various activator, repressor, and insulator elements. This capability is crucial, as it allows for capturing the complex dynamics among various regulatory elements, thereby offering a detailed insight into the potential effects of sequence replacement within a specific genomic locus (**Methods**).

In the case of the GATA1 locus, analysis of Enformer DNase predictions after replacing endogenous sequences with generated sequences showed that DNA-Diffusion sequences specific for K562 maintained or enhanced the predicted accessibility in the GATA1 enhancer for K562 compared to the native sequence, while showing no activity for GM12878 and HepG2, consistent with the ChromBPNet analysis (**Fig. 3d**). Similarly, when introducing cell type-specific HepG2 and GM12878 DNA-Diffusion sequences (**Fig. 3d**), we observed an increase in accessibility within previously non-activated GATA1 DHS regulatory regions in those cell lines. These sequences induced a statistically significant increase in accessibility in previously quiescent GATA1 DHS regulatory regions for GM12878 and HepG2, exceeding even the levels observed for the endogenous training sequences used during model training (t-test p-value < 0.001).

To assess the impact of these sequences on cell type-specific expression of GATA1, we analyzed the Enformer prediction of the CAGE (Cleavage and Polyadenylation Specificity Factor) assay within the 1KB region proximal to its TSS, noting that GATA1 is already expressed in K562 but not GM12878 and HepG2. The Enformer CAGE analysis revealed a significant increase in predicted promoter activity following the insertion of DNA-Diffusion sequences for each cell type (**Fig. 3e**). For example, GM12878 DNA-Diffusion sequences replacing the GATA1 regulatory region show a mean CAGE signal of 8.83 in GM12878, surpassing the activity of the best endogenous GM12878 DHS training sequences (8.77, t-test p < 0.001) in the same cell type. A similar reactivation was observed for HepG2 DNA-Diffusion sequences (8.33) in HepG2 while K562 DNA-Diffusion sequences maintained or slightly increased expression of GATA1 in K562 (8.71).

An example of replacing a cell type-specific sequence generated for HepG2 is shown **Fig. 3f** where K562 loses accessibility, GM12878 maintains closed chromatin and HepG2 shows an increase in chromatin accessibility. In addition, this also shows a specific reactivation of GATA1 in HepG2 but not in GM12878.

Importantly, similar patterns were observed at the CD19 and HNF4A loci (**Supplementary** Figs. 3b,c **and 4b,c**), reinforcing the reproducibility and cell type-specific regulatory potential of DNA-Diffusion sequences. These sequences consistently demonstrated accessibility levels on par with or higher than endogenous DHS regions and were also able to induce the intended cell type-specific expression of the target genes. This evidence supports the use of DNA-Diffusion synthetic sequences in regulatory circuits that may depend on diverse regulatory signals already present in a given genomic locus.

Taken together, these analyses highlight the model’s versatility in creating sequences with cell-specific potential in modulating chromatin accessibility and gene expression, showcasing its broader applicability and effectiveness in varied genomic and cellular contexts.

### Selection of optimal region for insertion of a synthetic sequence for reactivation of a target gene

To explore the effects of replacing DNA-Diffusion sequences beyond putative regulatory regions annotated based on known cell type-specific DHS sites, we aimed to pinpoint novel regions suitable for the insertion of synthetic elements that could enhance cell type-specific reactivation of a target gene, considering the potential interactions with other regulatory elements within the same locus.

We implemented a tiling strategy, analyzing every 200bp segment within the GATA1 locus (**Fig 4a**). By substituting a DNA-Diffusion sequence tailored for HepG2 into consecutive windows, we assessed changes in GATA1 expression, as reflected by the predicted CAGE signal around its TSS (**Fig 4b**).

**Figure 4.**
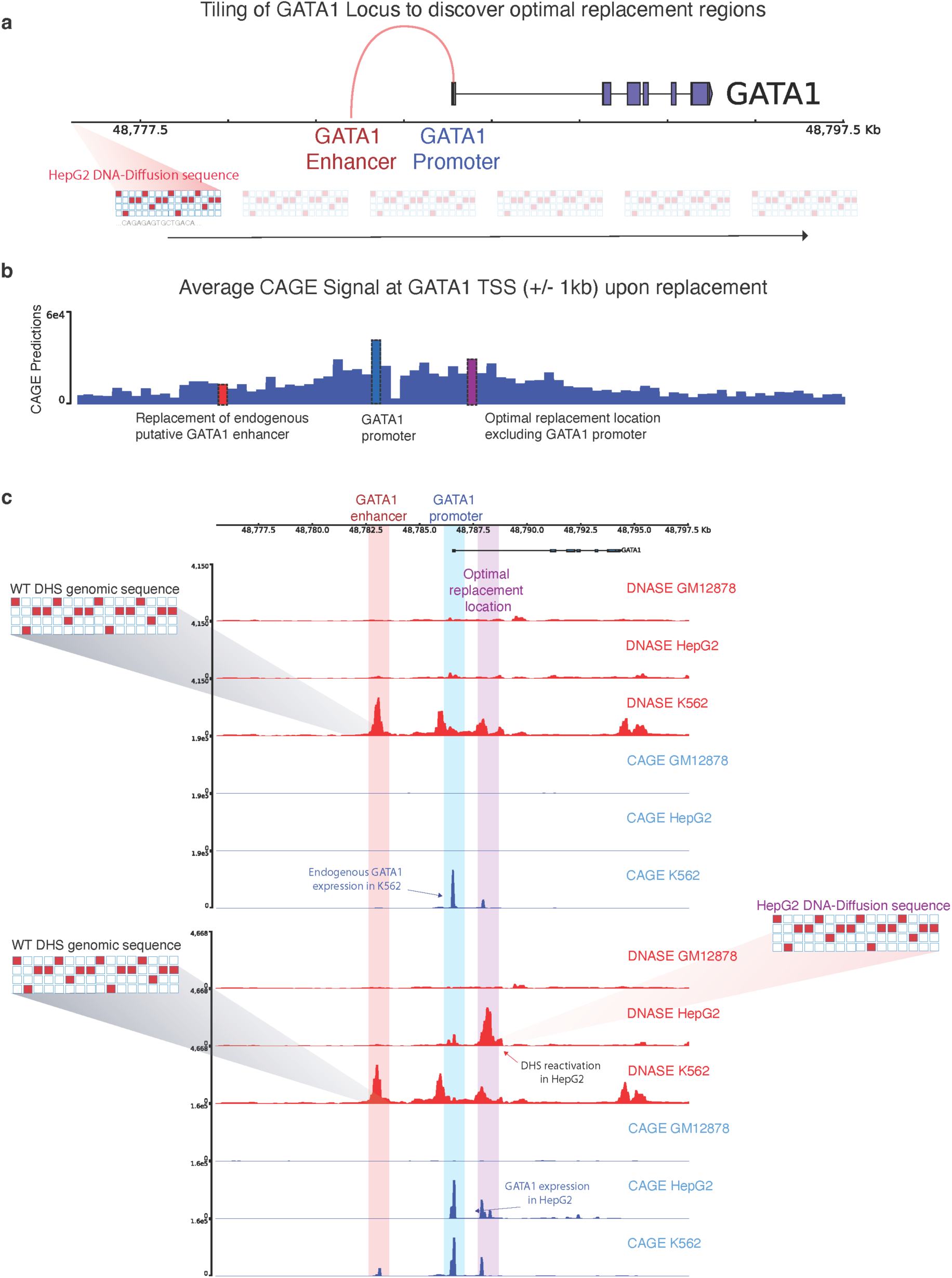
a) Tiling approach for exploring the optimal location of replacement of a synthetic sequence to reactivate a gene of interest. b) The average CAGE predictions for the GATA1 TSS +/− 1KB are shown for each replacement of the same HepG2-specific sequence throughout the entire locus. Key regions with strong predicted functional effects on GATA1 reactivation are highlighted; the original GATA1 enhancer in K562 (red), the GATA1 promoter (blue), and the optimal replacement region discovered by the tiling procedure (purple). c) Top: DNase and CAGE activity predictions at the GATA1 locus based on the wild-type sequence. Bottom: post-insertion of a HepG2-optimized DNA-Diffusion sequence into the optimal replacement location, resulting in heightened chromatin accessibility and GATA1 expression in HepG2, maintenance of accessibility and GATA expression in K562 and no significant changes in GM12878.

As already discussed in the previous section, substituting this sequence at the endogenous K562 DNase-accessible region substantially increased chromatin accessibility in HepG2 cells but also reactivated GATA1 expression (**Fig 3c**), here we investigated whether other locations might exhibit similar or improved reactivation effects. This analysis indicated that the level of predicted reactivation varied with the sequence replacement location. Interestingly, certain regions enhanced GATA1 reactivation in HepG2 cells beyond what was observed at the initial replacement location determined by the existing regulatory element in K562, specifically in an intronic region of GATA1, with the promoter region of GATA1 also exhibiting significant reactivation (**Fig 4c; Supplementary** Figure 8). Importantly, replacing the sequence outside an existing regulatory element for a given cell line offers a crucial advantage: it avoids disrupting the function of preexisting regulatory elements crucial for the cell type where the gene is naturally active.

This method effectively identified both optimal sites for element insertion and previously unrecognized regulatory regions, thereby providing a versatile approach for precise genomic modifications, considering the broader genomic context for any gene of interest. Nonetheless, it’s essential to recognize that our strategy, which relies on Enformer, may not fully account for the impact of regulatory elements distant from the TSS of the target gene, as suggested by prior studies analyzing the predicted effects of these models on the perturbation or deletion of distal regulatory elements^35^.

In summary, we demonstrated that DNA-Diffusion sequences can be introduced into previously uncharacterized regions, which upon replacement, may exhibit even a strong cell type-specific chromatin accessibility and gene reactivation as compared to annotated DHS sites. This capability marks an important step towards innovative regulatory engineering that does not depend solely on existing regulatory elements.

### Selection and characterization of synthetic sequences with different regulatory potential and cell type specificity

Having established that our DNA-Diffusion sequences match or exceed the accessibility and gene expression levels of the DHS sites used for training across different loci and cell types, we explored strategies for selecting sequences with particular regulatory characteristics along two dimensions: “signal intensity” and “signal specificity” (**Fig. 5a**). Generating cell type-specific sequences that fine-tune chromatin accessibility and gene expression can greatly enhance the development of synthetic gene circuits and precision gene therapies, potentially leading to innovative therapeutics for targeted drug release, diagnostic applications, or the dynamic modulation of cellular functions. Importantly, the generative aspect of our model enables us to sample a broad set of sequences to fulfill specific design requirements.

**Figure 5.**
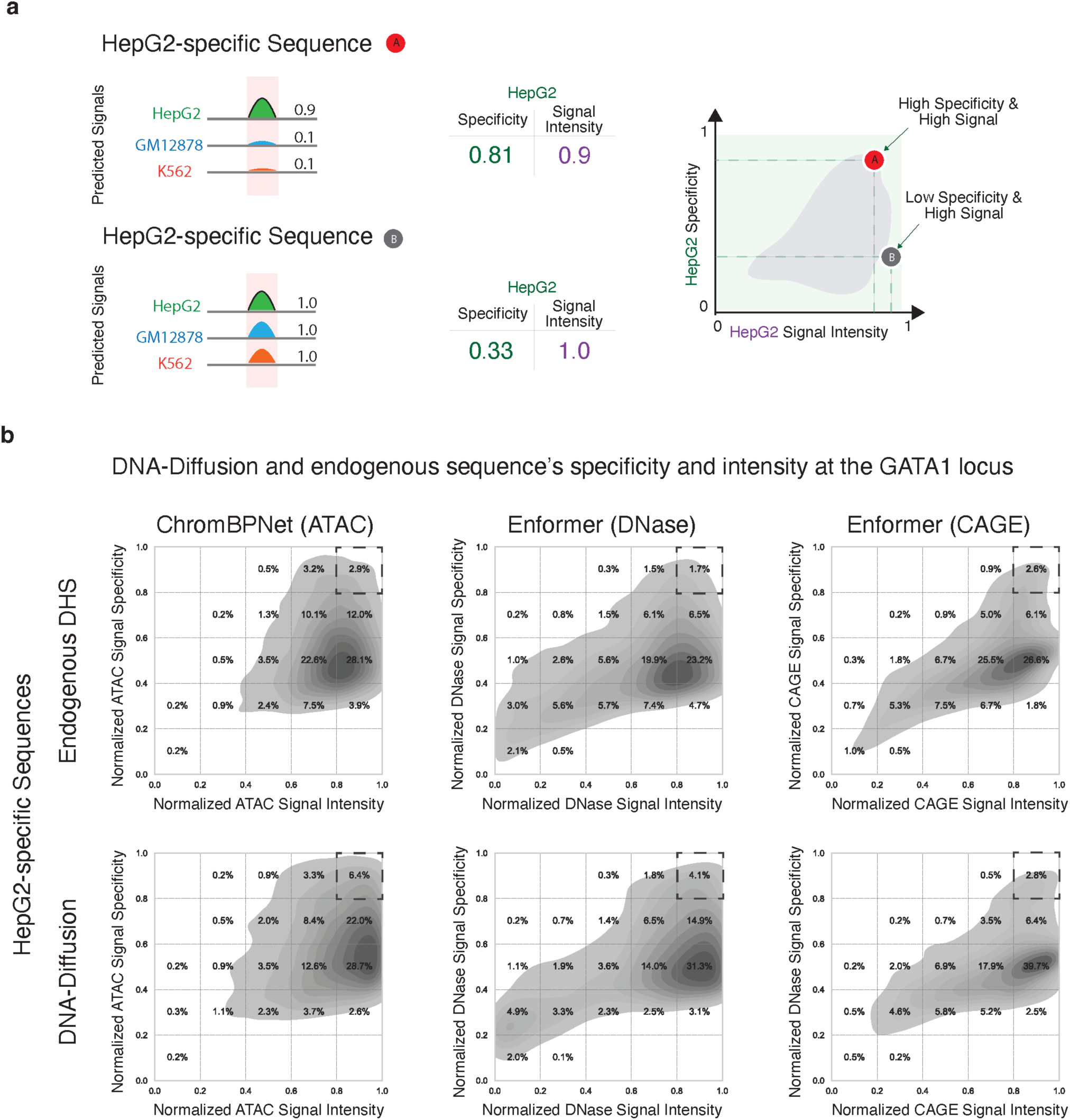
a) Schematic illustrating the workflow for evaluating both signal intensity and target cell specificity of endogenous and DNA-Diffusion sequences. Shown are two hypothetical sequences: A, a strong signal present in only one cell line (HepG2), and B, a strong signal present in all the cell lines (GM12878, HepG2, K562) b) Density plots depicting the distribution of signal intensity and specificity for both endogenous train DHS and DNA-Diffusion sequences. Dashed squares highlight the upper right corner (high signal, high specificity) of each plot, indicating the region of interest for the selection of potent and cell type-specific regulatory sequences.

To this end, we developed a framework to select DNA-Diffusion sequences that can display different levels of accessibility and regulatory activity across different cell types for a given locus of interest. Specifically, we combined *in-silico* oracles used to evaluate the regulatory potential of these sequences, averaging the Enformer CAGE, Enformer DNase, and ChromBPNet ATAC predictive scores to define overall signal specificity and specificity (**Methods**). Briefly, we characterized signal specificity for a particular target cell and sequence by calculating the ratio of the signal intensity in the target cell to the total signal intensities across all cells.

Comparing the training endogenous sequences with DNA-Diffusion sequences across these two metrics for the GATA1 locus, particularly focusing on HepG2-specific sequences, we noted significant variations. Despite the use of specificity filtration criteria during training, aimed at promoting cell type-specific sequences, the specificity of these sequences showed substantial variability, a trend effectively mirrored by the DNA-Diffusion sequences (**Fig 5b,c**). Remarkably, a larger number of DNA-Diffusion sequences exhibited superior performance, outperforming the endogenous sequences in both signal strength and specificity. This highlights the model’s refined capability to discern sequence features unique to specific cell types.

Following the observation of variations among DNA-Diffusion sequences, particularly with a subset showing outstanding performance in signal strength and specificity, we decided to investigate the defining features of these top-performing sequences further. We carried out a motif enrichment analysis to compare the sequences identified as high-signal with those deemed high-specificity **(Fig. 6a,b)**. This analysis aimed to identify overrepresented motifs within each subgroup to understand the sequence characteristics that contribute to their remarkable performance and cell type-specific effectiveness **(Fig. 6c)**.

**Figure 6.**
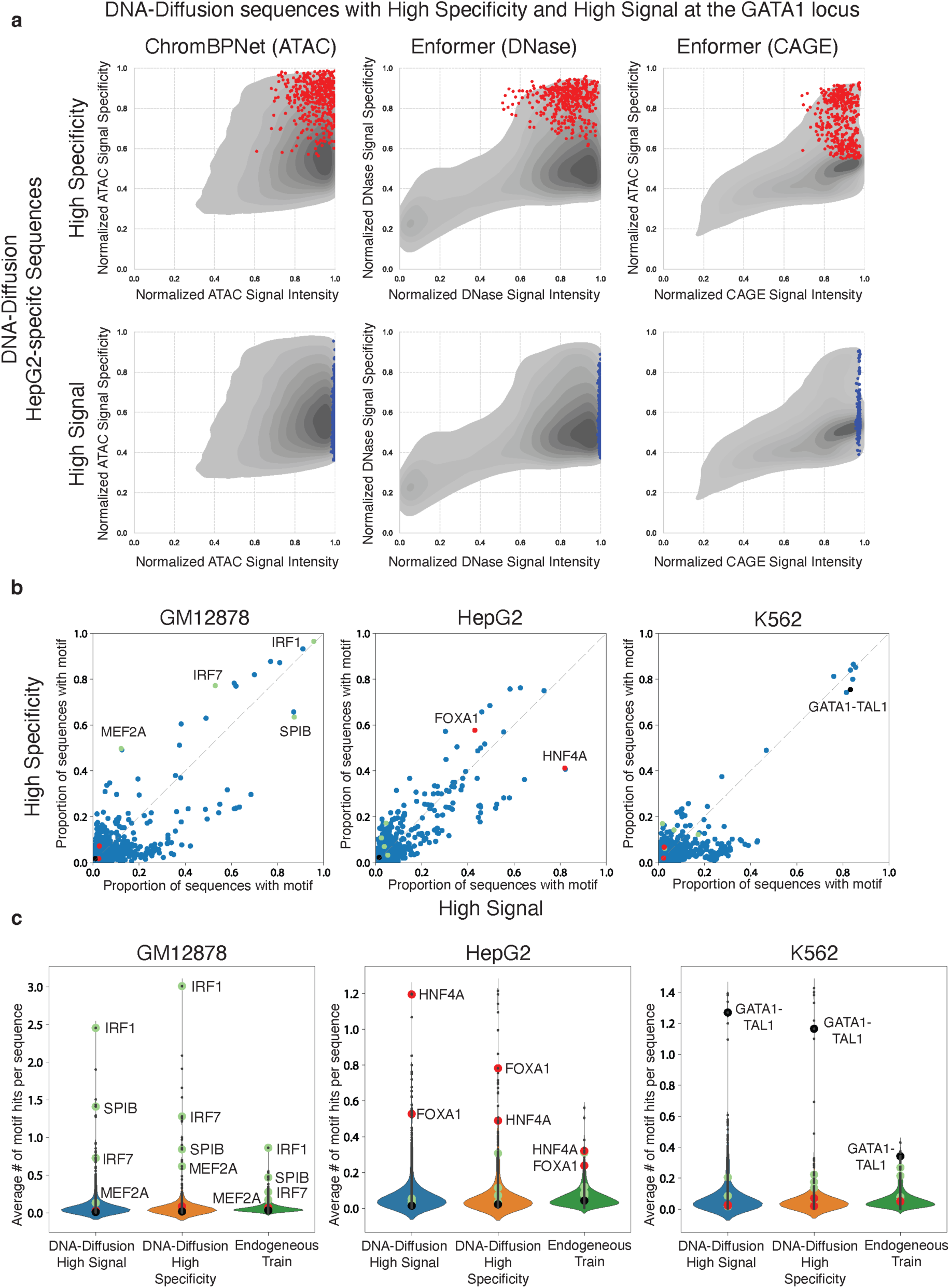
TF Motif Composition across High Signal and High Specificity DNA-Diffusion sequences. a) Selection of 400 high specificity (red) and 400 high signal (blue) HepG2-specific DNA-Diffusion sequences based on the activities at the GATA1 locus. Points represent these two sets of sequences and are overlayed on the density plot of 100k HepG2-specific sequence. b) The normalized proportion of sequences containing specific TF motifs is shown for the high signal and high specificity sets. Green, red, and black TF motif points highlight TFs known to be active in GM12878, HepG2, and K562 cells, respectively. c) Violin plots depict the average number of TF motif hits per sequence across high signal, high specificity, and endogenous train sets. Green, red, and black TF motif points are marked for literature-backed TFs for GM12878, HepG2, and K562 cell lines, respectively.

For GM12878 and HepG2, multiple TF motifs were predominantly observed in sequences classified as high-signal or high-specificity. Notably, in HepG2 sequences, a significant enrichment of the HNF4A motif was found in high-signal DNA-Diffusion sequences, with 82% showcasing this motif. In contrast, high-specificity sequences displayed the HNF4A motif less frequently, at 41%. Conversely, the FOXA1 motif was present in 58% of high-specificity sequences, compared to 48% in high-signal ones, suggesting HNF4A’s role in enhancing accessibility in HepG2, while FOXA1 might contribute to achieving greater specificity in this cell type.

In GM12878 sequences, immune-related TFs, such as MEF2A (50% in high-specificity vs. 12% in high-signal) and IRF7 (77% vs. 53%), were predominantly found in high-specificity sequences. Contrastingly, SPIB, another immune-associated factor, was more prevalent in high-signal sequences (64% vs. 87%). Additionally, IRF1 showed significant enrichment in both high-signal and high-specificity sequences (96% vs. 97%).

Interestingly, for K562 sequences, our analysis suggested fewer TFs (including GATA1-TAL1) are required for generating sequences with high specificity and signal intensity. However, our framework couldn’t distinguish between the two categories solely based on motif composition. The average motif hits for the aforementioned TFs also increased in their respective categories compared to others and endogenous training sequences. For instance, HNF4A had an average of 1.195 hits in high-signal DNA-Diffusion sequences, 0.49 hits in high-specificity sequences, and 0.32 hits in endogenous training sequences (**Fig. 6c**).

Further investigation of TF expression across cell types revealed that many TFs enriched in high-specific and high-signal sequences are predominantly expressed in a cell type-specific manner (**Supplementary** Fig. 9a). These findings suggest a sequence-based regulatory logic distinguishing these two categories of sequences, underscoring the nuanced relationship between TF motifs and their regulatory impact in specific cellular contexts.

## Discussion

Generative AI models have revolutionized various fields, including image, natural language processing and more recently biotechnology^36^. For example, DALL-E 2^37^, Chat-GPT^38^, and Stable Diffusion^9^ have shown great success in generating complex and realistic visual and textual content. Among these models, diffusion probabilistic models^7^ have shown great promise in various applications due to their ability to generate high-quality samples through a denoising process. Recently, in the realm of biotechnology, these generative models have shown remarkable success in protein synthesis ^39,40^ demonstrating their adaptability for tackling complex biological challenges. One of them is the generation of synthetic DNA sequences that could offer important downstream therapeutic applications.

In this study, we introduced DNA-Diffusion, a novel generative approach leveraging diffusion probabilistic models for the design of cell type-specific DNA regulatory sequences. Our findings demonstrate the model’s remarkable capacity to generate sequences that not only retain the essential properties of endogenous sequences, such as transcription factor binding motif composition, but also exhibit enhanced regulatory activities and accessibility specific to the targeted cell types.

Importantly, we propose a framework to guide and refine the search space for sequences with specific regulatory activity by optimizing their properties across different molecular features related to gene regulation (chromatin accessibility, expression level) and genomic location of insertion. This framework aims to fine-tune sequences for optimal performance in real-world applications, maximizing the potential of synthetic sequences to maintain their observed characteristics in practical scenarios.

DNA-Diffusion stands out from existing generative models in its ability to learn cell type-specific transcription factor grammar and generate sequences with enhanced accessibility and regulatory activities. Importantly, our model presents the first end-to-end solution to sequence design that requires no external model guidance nor orchestration of different models to achieve specificity. This contrasts with earlier methods like GANs and CNNs, which often require the coordination of multiple models and lack the end-to-end training capability across different cell types. In fact, while these models require complex layers and multiple models for different cell types, DNA-Diffusion offers a unified, end-to-end solution, streamlining the generation process.

While DNA-Diffusion marks a substantial advance, it is not without limitations. The current model relies on binarized labels for DHS peak identification, which might oversimplify the complexity of chromatin accessibility landscapes. Future iterations could benefit from incorporating continuous measures of chromatin accessibility to capture more nuanced regulatory dynamics. Additionally, expanding the model to encompass a broader range of cell types from the DHS index dataset would be an invaluable extension, allowing for more comprehensive applications in diverse biological contexts.

Additionally, exploring the model’s potential in disease contexts, such as cancer or genetic disorders, could open new therapeutic avenues. Specifically, generating regulatory elements that can modulate gene expression in these contexts could lead to novel, patient-specific treatments.

To translate in silico predictions of DNA-Diffusion sequences into tangible biological outcomes, robust experimental techniques that are able to replace endogenous sequences with generated sequences are paramount. These approaches would validate and elucidate the true functional potential and applicability of these designed sequences.

In conclusion, DNA-Diffusion exemplifies the potential of generative AI in advancing our understanding of genomic regulation and synthetic biology. By generating functional, cell-type-specific regulatory elements, this model holds immense promise for the future of precision medicine and synthetic biology, marking a significant milestone in the journey toward effective therapeutic strategies aimed at fine-tuning gene expression.

## Acknowledgments

We would like to acknowledge Bo Wang, Vallijah Subasri, Micaela Consens, Rex Ma, Judith Franziska Kribelbauer, Bart Deplancke, Jiecong Lin and other people in the Pinello Lab, Niccolò Zanichelli and the OpenBIOML community for helpful feedback or discussions. Anshul Kundaje, Anusri Pampari for their support in using the ChromBPnet models. L.P. is partially supported by 1R35HG010717-01. W.M. is partially supported by 1R35HG011317-01.

## Author Contributions

L.F.D.S, W.M., and L.P. conceptualized the project and its scope. L.F.D.S, Z.N., and S.S. designed, implemented and trained the diffusion model. Z.M.P., L.F.D.S, and S.S. performed the model interpretation and TF motif analyses. A.J.R and T.A.K conducted computational MPRA training and analysis. S.G, L.P., L.F.D.S, and S.S performed the ChromBPNet analysis. C.M.V.C, L.F.D.S, S.S, and Z.M.P conducted the BLAT analysis. A.W., T.M.T, Z.L, L.F.D.S, and S.S. assisted with Enformer analyses. M.M, N.W., C.C., W.C., C.S., and M.B. assisted with model testing and exploration. P.C., Z.L, and L.F.D.S curated the control datasets. L.F.D.S, S.S., Z.M.P, A.J.R, S.G, Z.N. A.W, N.W, T.T, M.M, and V.S., B.W., W.M., and L.P assisted with manuscript writing. L.F.D.S, S.S., Z.M.P, and L.P prepared the manuscript and associated figures with input from all authors.

## Methods

### Data preprocessing for training

To facilitate data filtering and analysis, the binary peaks annotation from the DHS Index was augmented with the following metadata columns: chromosome (chr), DHS start position, DHS end position, DHS width, DHS summit (peak maximum), total signal strength, component, proportion, and sequence ^19^. Given the variable length of ENCODE DHS sites, we standardized sequence analysis by extracting a 200bp region centered on the peak summit (+/− 100bp). We use this threshold by assuming that the peak center should represent the most accessible regions and take into consideration the expected number of nucleotides protected by a single chromosome ^18^. Our analysis of 733 biosamples from 438 cell types aim to identify replicates exhibiting both strong component association in the non-negative factorization data and ease of laboratory culture; ultimately yielding four replicates from K562, GM12878, HepG2, and hESCT0 cell lines, respectively.

Using the selected replicates, the complete dataset was filtered only to include the DHS sites that were cell-type specific. This cell type-specificity refinement strategy consisted of prioritizing peaks that had limited occurrences in cell lines outside our replicates, while also putting emphasis on samples that had peaks in other replicates of the same cell. From there we filtered out samples that occurred in more than one of our cells of interest, so that we were left with a dataset that was sorted in descending order of peak presence in other replicates and ascending order for peak presence in other cell types. GM12878 was the cell line with the smallest number of cell type-specific peaks (11968 sites). To have a balanced number of regions across all cells every replicate was restricted to the same number. From there, regions present in chr1 and chr2 were held out as test and validation shuffle sets, respectively.

### Model Architecture

The DNA-Diffusion model takes inspiration from the Annotated Diffusion Model ^41^, with some key modifications. At its core, the diffusion model consists of a U-Net ^17^ that first projects the batch of encoded sequences to the desired channel dimension of 200. Each downsampling layer consists of 2 ResNet ^17,42^ blocks, followed by a linear attention ^43^ layer, and then downsampling convolutional layers until the output channel dimension is reached. The middle layers of the U-Net consist of a ResNet block and an attention layer before an additional ResNet block. The structure of the upsampling stage is the same as the downsampling stage with the exception of the downsampling convolutional layers being replaced with a upsampling convolutional layers. The output of the U-Net is then projected through one final ResNet block before one convolutional layer is applied to return the output with the same dimensionality as the input.

During the forward process of this diffusion model, a batch of DNA sequences and their corresponding cell type-specific labels are fed through the forward process where the cell type labels are randomly masked (setting to null), allowing for classifier-free guidance^44^ during training and downstream sampling. Using this sampling process 100,000 sequences per cell type were generated, resulting in a generated set of 300,000 sequences.

### Training process

The main diffusion model was implemented in PyTorch 2.0.0 and trained on 4x 40gb Nvidia A100s using a batch size of 960 for 2000 epochs. For distributed computing across the nodes we employed Slurm Workload Manager. We used the Adam optimizer (reference) with a learning rate of 1e-4 along with PyTorch default values for other hyperparameters: *β*_1_ = 0.9 and *β*_2_ = 0.999. Linear noising schedule was used with *β_start_* = 0.0001 and *β_end_* = 0.005.

### In-Silico Validation

#### Assessing DNA chromatin accessibility with ChromBPNet

Accessible chromatin is a notable marker of regulatory activity for a DNA sequence. We use the ChromBPNet model as a biological oracle to assess the cell-type specificity of generated diffusion sequences’ chromatin accessibility. ChromBPNet consists of two convolutional neural networks that are similar in structure to BPNet^46^. One convolutional neural network learns to predict base-pair resolution chromatin accessibility, while the second network learns to predict the noise of an experimental assay (e.g. DNase-Seq, ATAC-seq). The outputs of the two networks are summed to predict the log of total counts and base-resolution probability distribution of counts of a 1000 bp sequence. Because the ChromBPNet architecture consists of two models to predict the ground truth data of an assay, the first convolutional neural network’s output can serve as a bias-corrected chromatin accessibility at base-pair resolution without the noise of an assay affecting the prediction.

We used ChromBPNet models trained on ENCSR000EPC, ENCSR000EMT, and ENCSR000ENP to predict base-pair chromatin accessibility of DNA-Diffusion sequences in K562, GM12878, and HepG2 contexts respectively. Each ChromBPNet model’s input was a one-hot encoded 2114 bp DNA sequence (A = [1,0,0,0], C = [0,1,0,0], G = [0,0,1,0], T = [0,0,0,1]).

To evaluate how DNA sequences generated by the diffusion model affect chromatin accessibility, we inserted them within known or putative regulatory regions for the following genes: GATA1, CD19, and HNF4A and calculated the mean of the predicted base-resolution chromatin accessibility score.

Given a generated sequence designed for a particular cell type, we ideally expect that the ChromBPNet model trained to predict accessibility in that cell type will yield greater chromatin accessibility predictions relative to the other two ChromBPNet models, indicating greater regulatory potential. However, the three ChromBPNet models used were trained on experimental data with different read depths: ENCSR000EPC had a read depth of 245 million, ENCSR000EMT had a read depth of 68 million, and ENCSR000EMT had a read depth of 323 million. Thus, we observed that models trained on deeper data typically predicted higher chromatin accessibility compared to the other two models for a generated sequence, irrespective of the cell type specificity of a sequence – leading to inaccurate comparison of model outputs. To combat this potential confounder, we applied quantile normalization (see **Normalization and scaling of the data**).

#### MPRA Predictor

We train our MPRA activity predictor using MPRA data from GM12878, K562, HepG2, SK-N-SH, and A549 cells (accession ids are available in **Supplementary Table 3**), collected by one lab using a uniform protocol. The assayed sequences are 200bp long and most of them are derived from human genomic segments containing the reference and alternate alleles for variants from UK Biobank and GTEx. These sequences are cloned upstream of a reporter gene and this construct is delivered to host cells using transient transfection. Each sequence’s MPRA activity is measured as the log2 of the ratio of the number of mRNA molecules produced using the construct to the number of constructs delivered to the host cells. From ENCODE, we obtain 318734, 636185, 750298, 750084, and 318734 MPRA activity measurements from GM12878, K562, HepG2, SK-N-SH, and A549 cells respectively.

To predict a given sequence’s activity in each of the cells, we input the 200bp sequence to Enformer and obtain its sequence embeddings from the last transformer layer. Then, we perform attention pooling (over the sequence’s length) to get the final sequence embedding. This sequence embedding is supplied to a fully connected output layer that simultaneously predicts MPRA activity in all five cell lines using multi-task learning. We initialize Enformer using pretrained weights provided by the authors. Following recent work from the lab that collected and used a portion of this data^11^, we trained our model using all sequences except those from chromosomes 7, 13, 19, 21, and X (∼79% of all sequences). Sequences from chromosomes 7 and 13 are used in the validation set (∼13% of all sequences), and sequences from chromosomes 19, 21, and X (∼7% of all sequences) are used in the test set. The model is trained for a maximum of 50 epochs using the AdamW optimizer^45^ with a 1e-4 learning rate and 1e-4 weight decay. Training is stopped if model performance on the validation set does not improve for 5 consecutive epochs.

Our trained model is quite accurate and achieves Spearman’s rank correlation coefficients of 0.4527, 0.7285, 0.7682, 0.7745 and 0.6682 on the test sets for GM12878, K562, HepG2, SK-N-SH, and A549 cells respectively. It achieves Pearson’s correlation coefficients of 0.4479, 0.8115, 0.8257, 0.8199 and 0.7218 on the test sets for GM12878, K562, HepG2, SK-N-SH, and A549 cells respectively.

We get model predictions for the generated sequences, control sequences, and sequences used in the training, validation, and test sets of the diffusion models. Since we do not have MPRA data from hESCT0 cells, we only analyze generated sequences that are designed to have regulatory activity in GM1278, K562, or HepG2 cells.

#### Assessing DNA chromatin accessibility and gene expression with Enformer

We replaced the endogenous sequence of putative regulatory regions with DNA-Diffusion or other endogenous sequences and considered Enformer predictions using a window of 393,216 base pairs around the region used for replacement. The resulting Enformer predictions offered an approximate 128 bp resolution window for DNase and CAGE tracks across all three cell lines. The extracted predictions across enhancer and promoter regions were averaged to generate a single value that described the prediction signal at the given coordinate.

A shortcoming of Enformer is that it is difficult to decrease or remove the CAGE signal in gene regions where a cell is already expected to display promoter activity. As a result, *in-silico* validation results were only utilized from regions where the cell line previously had no CAGE activity. This resulted in the following process: K562 sequence activity was examined in HNF4A and CD19, HepG2 in GATA1 and CD19, and GM12878 in HNF4A and GATA1 (**Supplementary Table 2**).

### Normalization and scaling of the data

Since Enformer (DNase and CAGE) and ChromBPNet (ATAC-seq) were trained using data with different sequencing depths, their baseline predictions exhibited a wide dynamic range, making it difficult to compare the difference in raw predictions across cell lines. Utilizing quantile normalization, we adjusted these predictions and rescaled the data to a common range. This process helped to minimize the differences and biases, allowing for more accurate comparisons and interpretations of the data across the loci and cell types.

For ChromBPNet predictions in all loci, we normalized the three cell lines’ accessibility predictions, generating a similarly distributed set across K562, HepG2, and GM12878. Instead of different models, Enformer outputs a separate prediction signal track for each cell type and data modality; we used the by-cell procedure mentioned from ChromBPNet to normalize the Enformer DNase and CAGE track predictions.

During normalization, we artificially introduced prediction values from 20k annotated GENCODE v43 protein-coding transcript promoters and 5k random endogenous genomic regions to ensure a final dynamic range representing natural endogenous sequence occurrence distribution.

During the ChromBPNet and Enformer DNase normalization, we introduced 5k prediction values captured from sequences selected directly from the random genome coordinates. The sequence coordinates were defined using the Bedtools Random function ^47^ in the hg38 genome version. While for the Enformer CAGE prediction normalization, we added prediction values from 20k promoters, utilizing predictions captured from their original genomic occurrence.

After normalizing all the signal predictions, we used the rank percentile to transform all the normalized values from ChromBPNet, DNase, and CAGE along each cell type to accommodate the values between a 0-1 scale.

### Cell type-specificity and ranking of sequences based on the predictions of different oracles

We call a predicted signal cell-specific when a prediction presents a bias towards a single cell type. For example, a GM12878 endogenous positive sequence is expected to have a stronger accessibility signal or targeted gene expression (e.g. GATA1) for GM12878 cells while presenting almost no signal in HepG2 and K562 cells.

We calculated the cell specificity for a given metric by selecting a specific cell type as the target and the other two remaining cells as off-targets. The specificity was calculated by dividing the signal in the target cell by the sum of the signal in all different cells (target + off-target1 + off-target2). The signal specificity is 1.0 when only the target cell shows activity. The signal specificity is 0.33 when all the cell types have exactly the same signal intensity.

We calculated specificity metrics for all the model predictions from ChromBPNet, Enformer DNase, and Enformer CAGE resulting in a total of twenty-four metrics per sequence and considered the average rank across these metrics as described below.

### DNA Diffusion Specificity Focused sequences

To evaluate the role of cell specificity in our model, we first filtered the set of sequences to keep a moderate baseline signal for all metrics (DNase, ChromBPNet, and CAGE > predicted signal percentile 0.5) and a baseline specificity of ∼0.5. We removed sequences that performed well only in a single metric to ensure a moderate baseline signal and specificity. After filtering out sequences with prediction signal and specificity under the baseline thresholds, we selected sequences presenting high specificity (> 0.8) in at least one of the metrics (DNase, ChromBPNet, and CAGE).

Sequences with high specificity in at least one of the metrics were ranked by creating first an averaged accessibility rank (DNase + ChromBPNet) and selecting the final sequences by averaging the previously generated accessibility rank and the CAGE rank.

DNase and ChromBPNet were ranked in their respective groups before being averaged to create an average accessibility rank. From there, the CAGE values were ranked and then averaged with the accessibility rank to obtain an overall rank across the baseline signal metrics. This two-step procedure allowed us to avoid the specificity being biased through accessibility since Enformer DNase predictions and ChromBPNet are correlated. This new rank was then utilized in conjunction with the designated high-specificity sequences to select a set of 400 highest-ranked DNA-Diffusion specificity-focused sequences.

### DNA Diffusion Signal Focused sequences

A different DNA-Diffusion sequence group was selected to investigate the role of signal strength without considering cell specificity. We selected 400 DNA-diffusion High Signal sequences for this set by ranking the strongest signal and averaging the rank for all metrics (DNase, ChromBPNet, and CAGE) in both cell-specific loci. The final top 400 sequences present the highest possible signal average among all generated sequences. No specificity metric was used in this group.

### Motif Scanning

To scan motifs the MOODS package was used using the vertebrate motifs from the JASPAR database (n = 949 PWMs). The parameters used in MOODS were a p-value of 0.0001 and a pseudocount (--ps 0.0001).

**Supplementary Table 1.** Cell type specific genes and regulatory regions selected for each cell type.

| Gene | Enhancer Region | Gene Region | Cell Type |
| --- | --- | --- | --- |
| GATA1 | chrX:48782929-48783129 | chrX:48785536-48787536 | K562 |
| HNF4A | chr20:44370692-44370892 | chr20:44355699-44432845 | HepG2 |
| CD19 | chr16:28930777-28930977 | chr16:28931971-28939342 | GM12878 |

**Supplementary Table 2.** Genes tested for reactivation for each cell type.

Genes tested for reactivation for each cell type
|  |  |
| --- | --- |
| **GM12878** | GATA1 |
|  | HNF4A |
| **K562** | HNF4A |
|  | CD19 |
| **HepG2** | CD19 |
|  | GATA1 |

**Supplementary Table 3.** ENCODE IDs of samples used for the training of the MPRA predictor model.

ENCODE IDs of samples used for the training of the MPRA predictor model
|  |
| --- |
| ENCFF996ECA |
| ENCFF018AMJ |
| ENCFF345ASG |
| ENCFF970OLE |
| ENCFF318XMJ |
| ENCFF821XQZ |
| ENCFF358MBK |
| ENCFF379XWL |
| ENCFF774CHX |
| ENCFF138DJM |
| ENCFF277DDE |
| ENCFF334EKU |
| ENCFF857FQR |
| ENCFF259NMG |
| ENCFF477LDL |
| ENCFF484JFE |
| ENCFF227KRF |
| ENCFF102ZVT |
| ENCFF418GRL |
| ENCFF333BAD |
| ENCFF307HBZ |
| ENCFF771HPB |
| ENCFF359KJL |
| ENCFF035HKU |
| ENCFF759PPO |
| ENCFF705AES |
| ENCFF256WKS |
| ENCFF352JAC |
| ENCFF147SMK |
| ENCFF311DJW |
| ENCFF350IJA |
| ENCFF815ORW |
| ENCFF402GOL |
| ENCFF865LNO |
| ENCFF755GRH |
| ENCFF440YVF |
| ENCFF703OIL |
| ENCFF927USI |
| ENCFF476FXK |
| ENCFF742ENC |
| ENCFF112HAT |
| ENCFF792IHA |
| ENCFF267VJ |

---

**Supplementary Figure 1.**
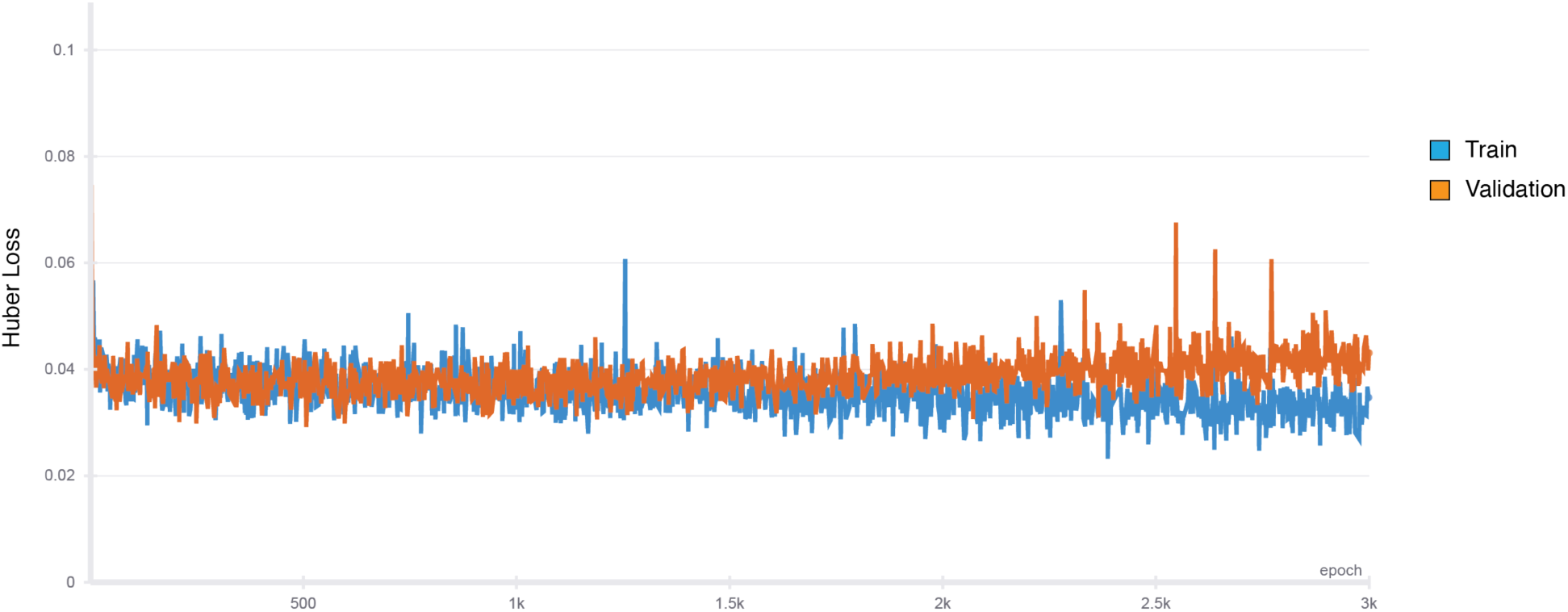
DNA-Diffusion training loss plot.

**Supplementary Figure 2.**
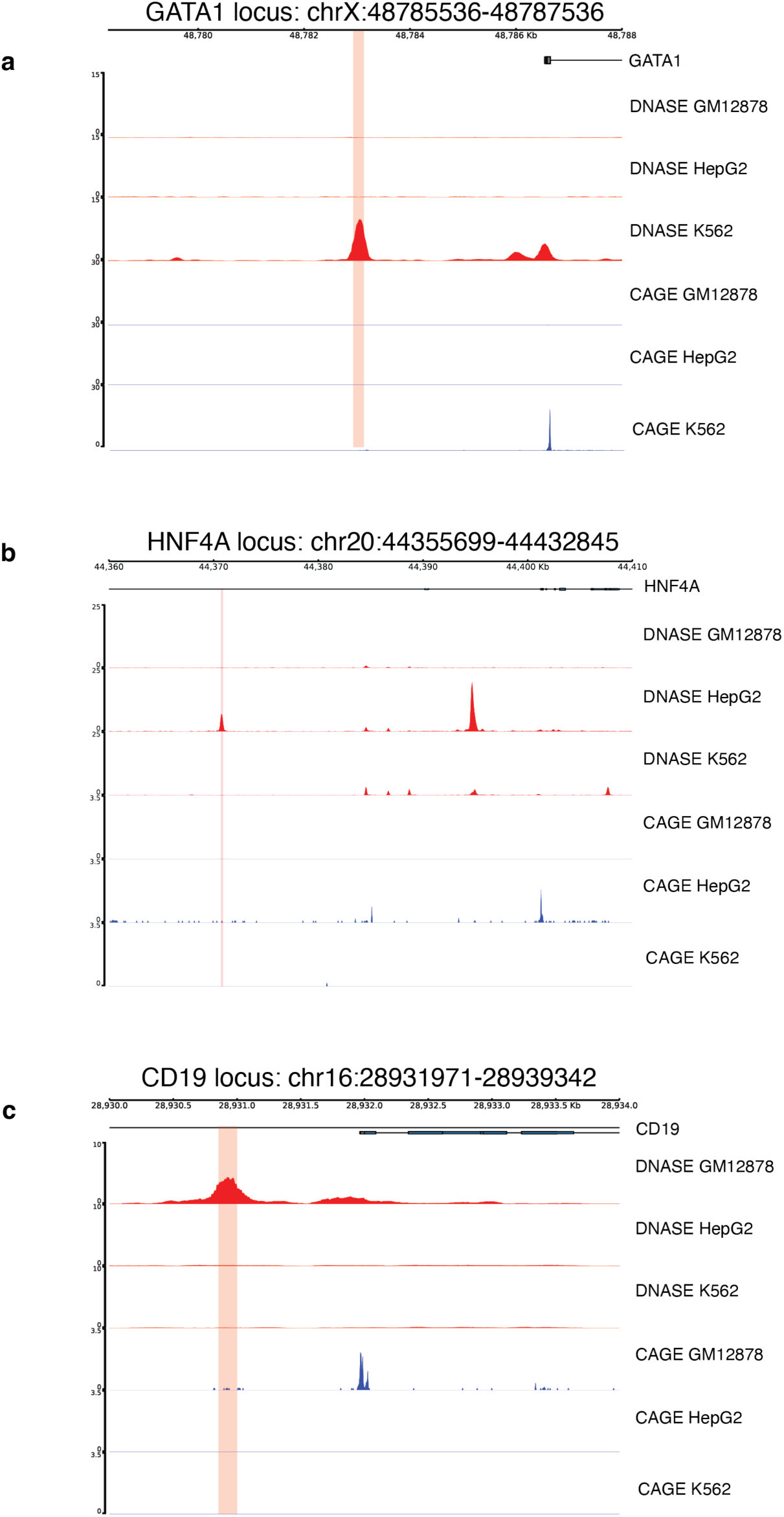
Endogenous DNase and CAGE Tracks. Chromatin accessibility (ENCODE DNase, red track) and gene expression (FANTOM CAGE, blue track) profiles are shown for three key gene loci. Each panel displays a window in the genomic region potentially controlling a specific gene (GATA1, HNF4A, CD19).

**Supplementary Figure 3.**
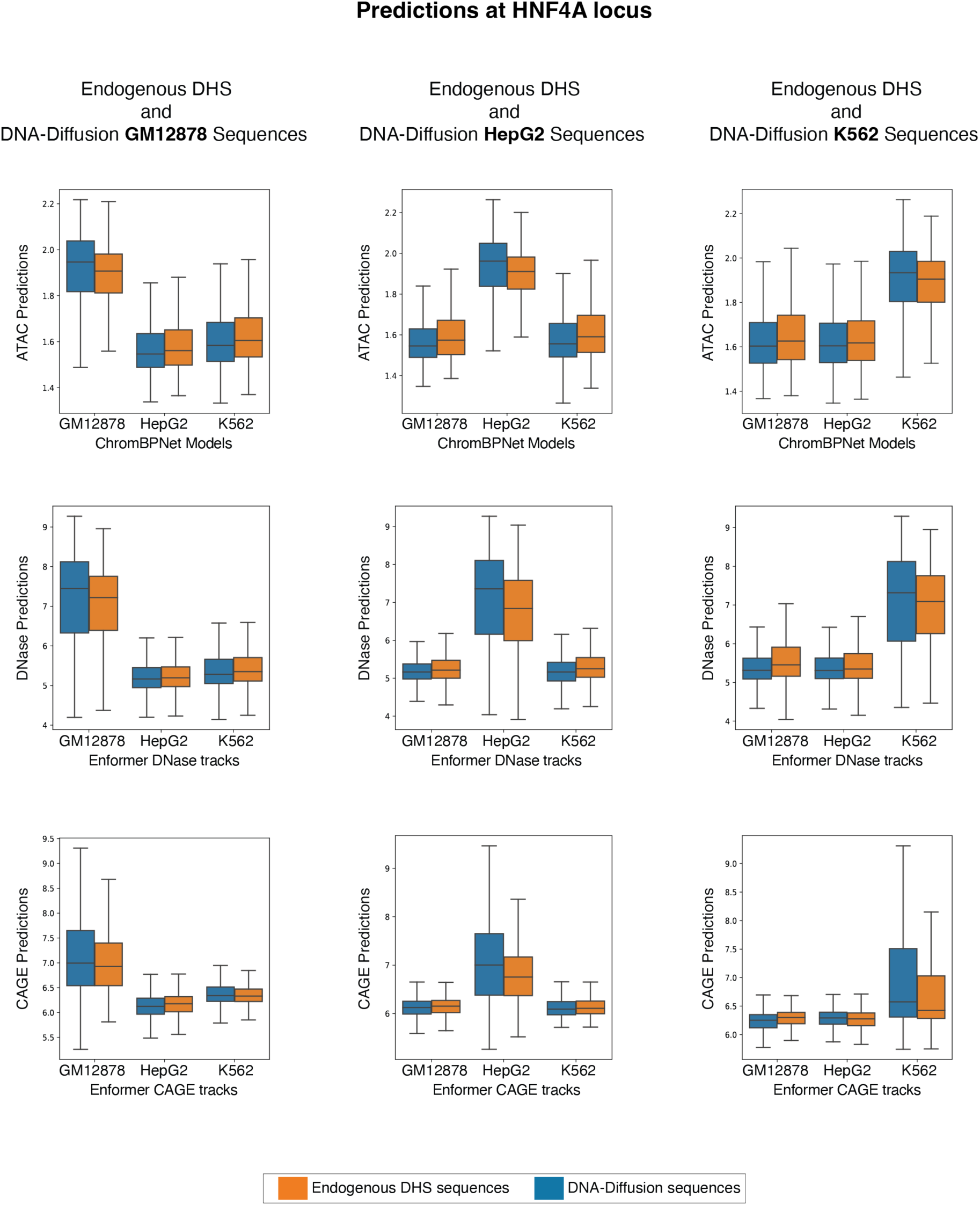
Endogenous train and DNA-Diffusion predictions across all three downstream oracles within the HNF4A gene locus. a) Boxplots showing the predicted chromatin accessibility/gene expression activity (ChromBPNet ATAC, Enformer DNase, Enformer CAGE) upon replacement within the HNF4A locus with endogenous DHS train and DNA-Diffusion sequences specific for each cell line (GM12878, HepG2, K562).

**Supplementary Figure 4.**
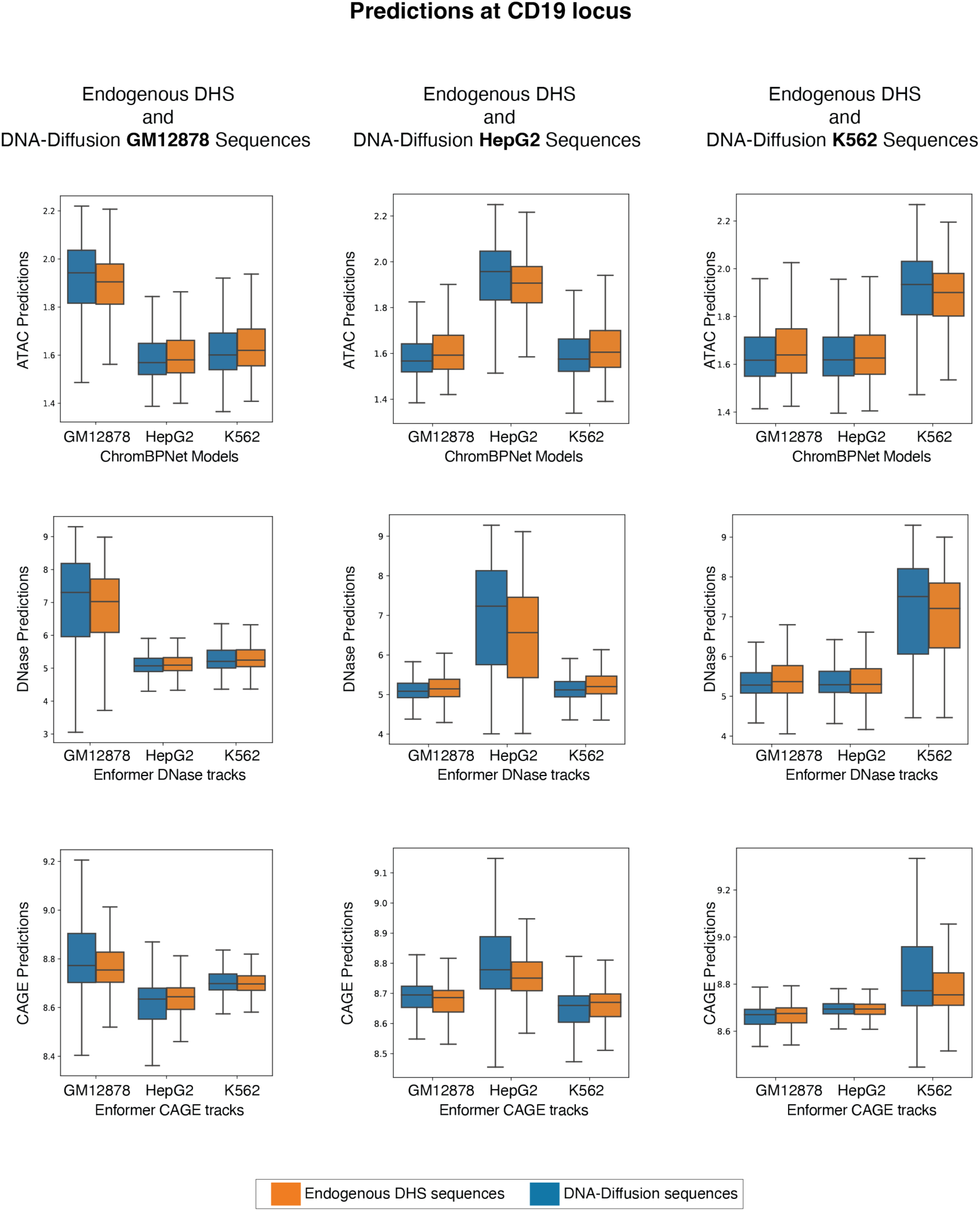
Endogenous train and DNA-Diffusion predictions across all three downstream oracles within the CD19 gene locus. a) Boxplots showing the predicted chromatin accessibility/gene expression activity (ChromBPNet ATAC, Enformer DNase, Enformer CAGE) upon replacement within the CD19 locus with endogenous DHS train and DNA-Diffusion sequences specific for each cell line (GM12878, HepG2, K562).

**Supplementary Figure 5.**
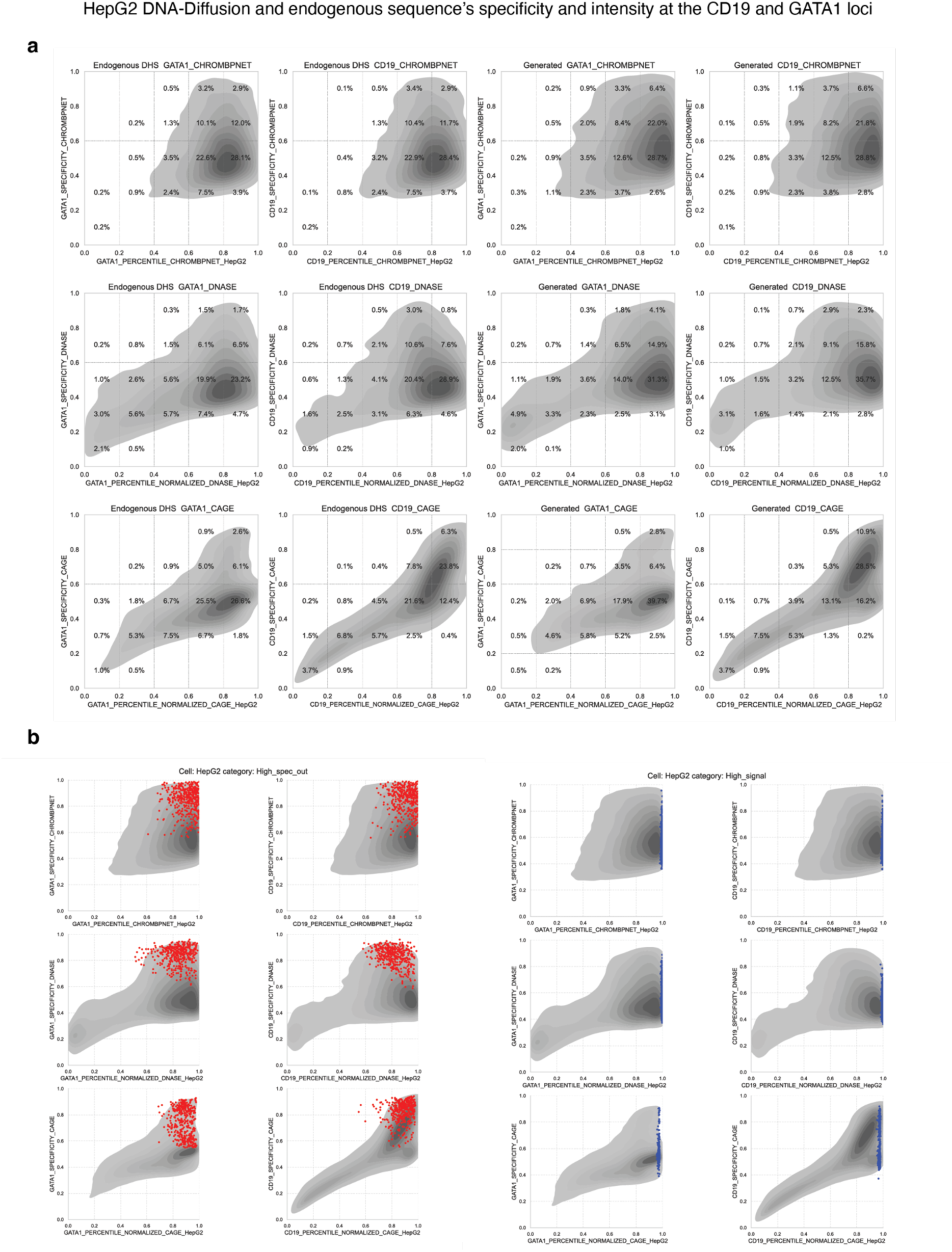
Oracle prediction scores with HepG2 cell type-specific sequences across CD19 and GATA1 gene loci.

**Supplementary Figure 6.**
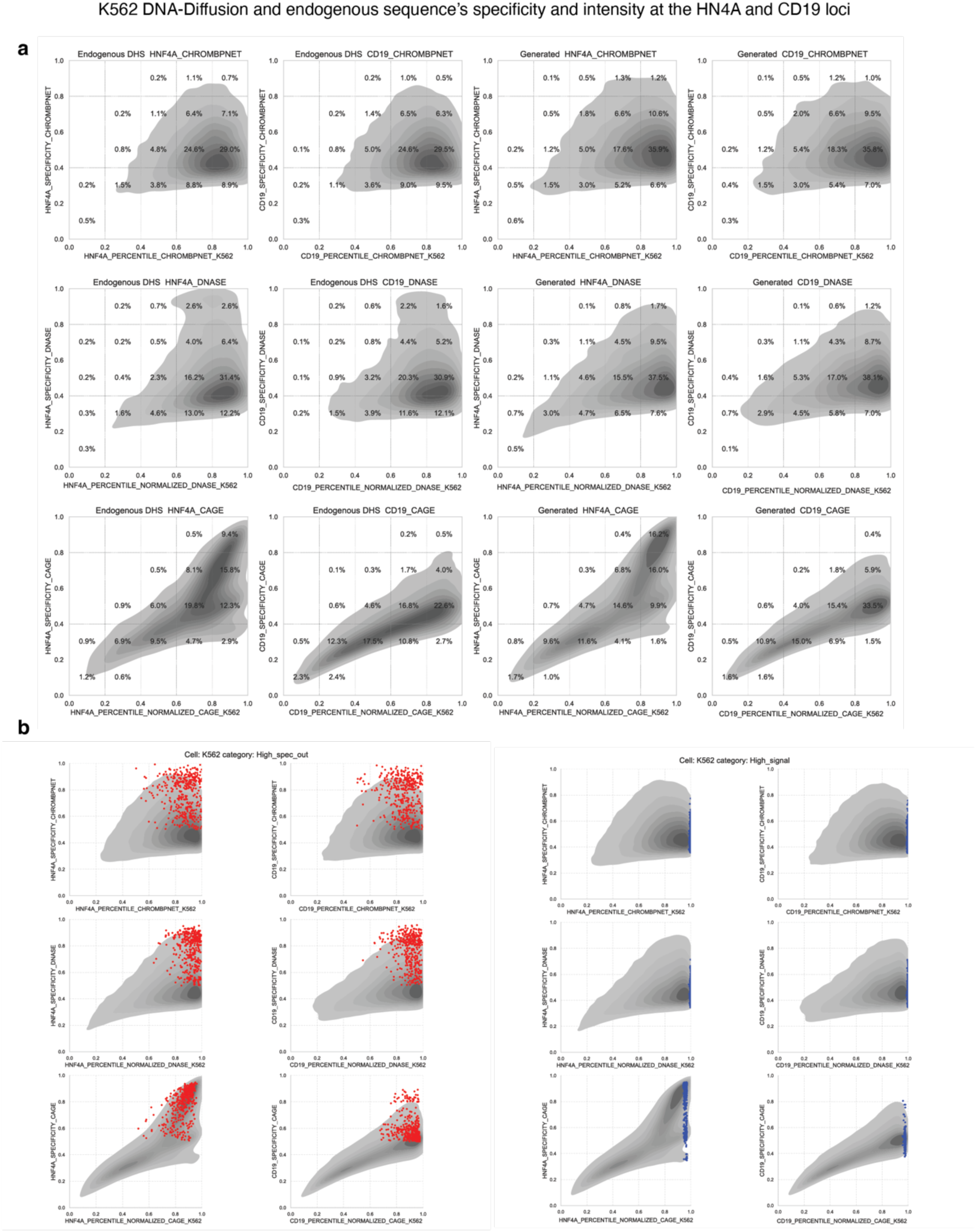
Oracle prediction scores with K562 cell type-specific sequences across CD19 and HNF4A gene loci.

**Supplementary Figure 7.**
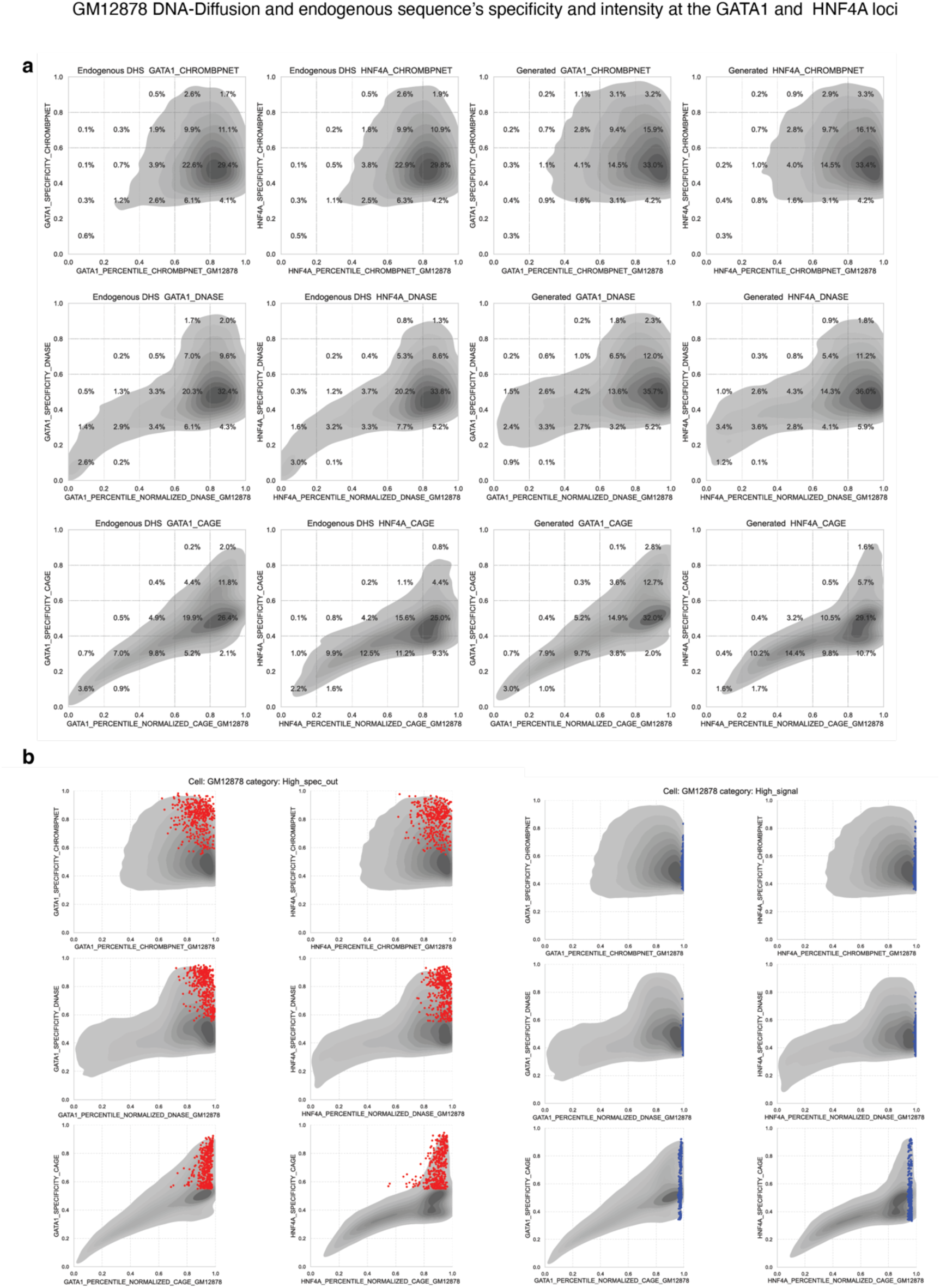
Oracle prediction scores with K562 cell type-specific sequences across HNF4A and GATA1 gene loci.

**Supplementary Figure 8.**
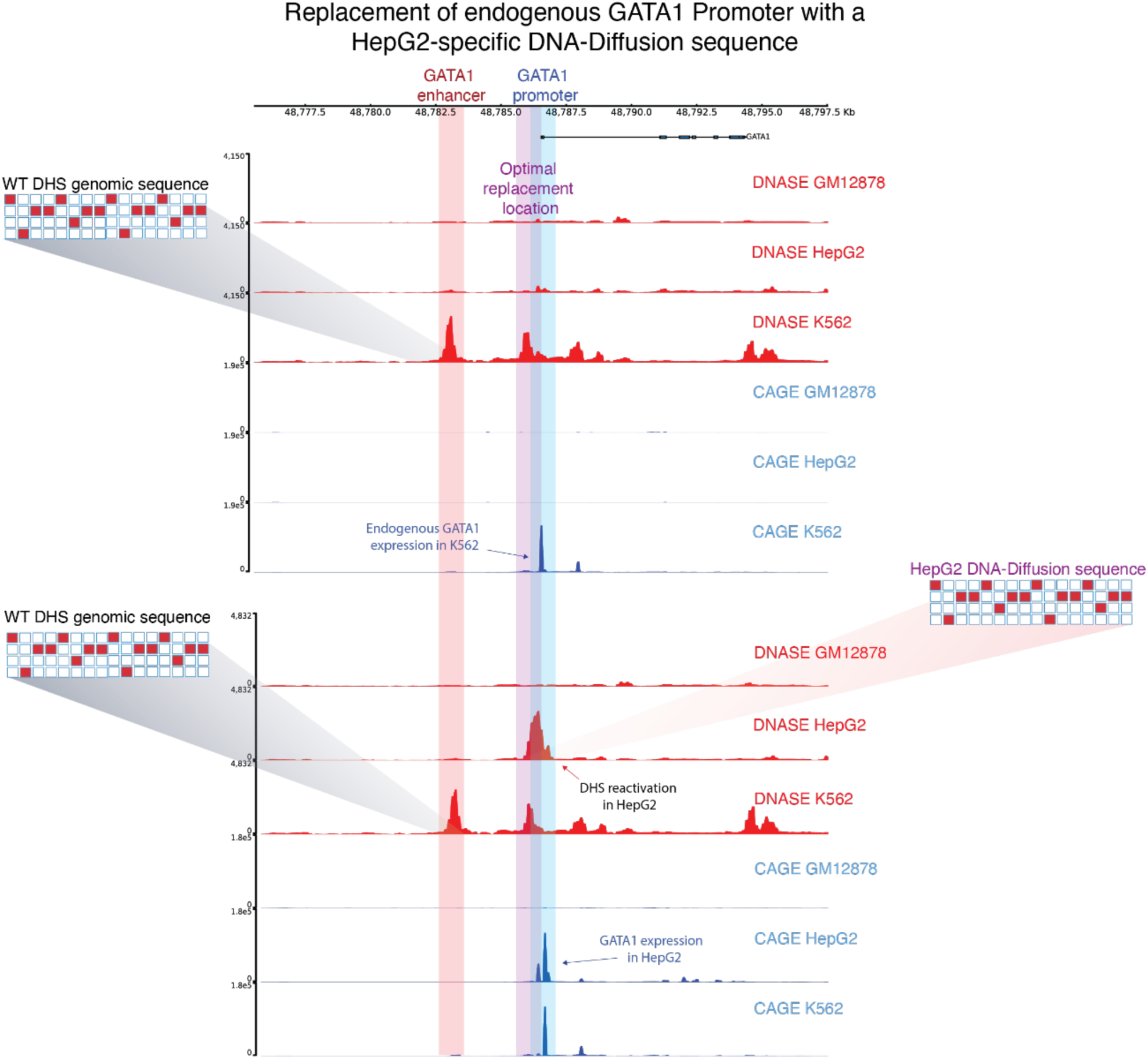
Exploring the effect of GATA1 promoter region replacement with HepG2-specific DNA-Diffusion sequence.

**Supplementary Figure 9.**
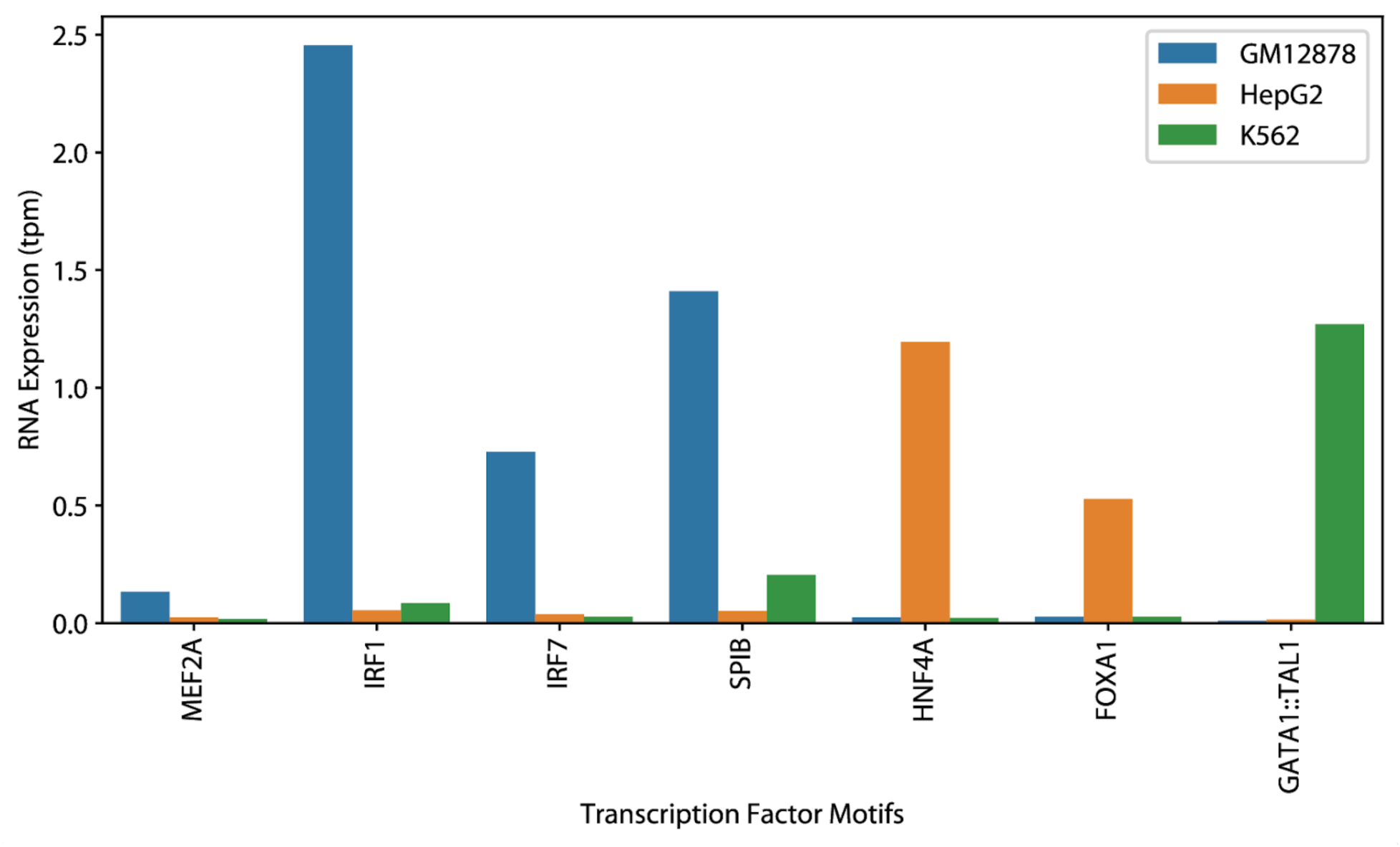
Endogenous RNA expression values for selected cell type-specific TF motifs. Bar plots showing TPM values for cell type specific TFs from **Fig. 6** across the three cell types.

## References

1. Kazachenka, A. et al. Identification, Characterization, and Heritability of Murine Metastable Epialleles: Implications for Non-genetic Inheritance. Cell 175, 1717 (2018).

2. Inoue, F. et al. A systematic comparison reveals substantial differences in chromosomal versus episomal encoding of enhancer activity. Genome Res. 27, 38–52 (2017).

3. Kundaje, A. et al. Integrative analysis of 111 reference human epigenomes. Nature 518, 317–330 (2015).

4. Martens, J. H. A. & Stunnenberg, H. G. BLUEPRINT: mapping human blood cell epigenomes. Haematologica 98, 1487–1489 (2013).

5. Noguchi, S. et al. FANTOM5 CAGE profiles of human and mouse samples. Scientific Data 4, 1–10 (2017).

6. Song, J., Meng, C. & Ermon, S. Denoising Diffusion Implicit Models. arXiv [cs.LG] (2020).

7. Ho, J., Jain, A. & Abbeel, P. Denoising Diffusion Probabilistic Models. Adv. Neural Inf. Process. Syst. 33, 6840–6851 (2020).

8. Li, X. L., Thickstun, J., Gulrajani, I., Liang, P. & Hashimoto, T. B. Diffusion-LM Improves Controllable Text Generation. arXiv [cs.CL] 4328–4343 (2022).

9. Rombach, R., Blattmann, A., Lorenz, D., Esser, P. & Ommer, B. High-resolution image synthesis with latent diffusion models. arXiv [cs.CV] 10684–10695 (2021).

10. Watson, J. L. et al. De novo design of protein structure and function with RFdiffusion. Nature 620, 1089– 1100 (2023).

11. Gosai, S. J., et al. Machine-guided design of synthetic cell type-specific cis-regulatory elements. bioRxiv (2023) doi:10.1101/2023.08.08.552077.

12. Taskiran, I. I. et al. Cell type directed design of synthetic enhancers. Nature 1–3 (2023).

13. Zrimec, J. et al. Controlling gene expression with deep generative design of regulatory DNA. Nat. Commun. 13, 5099 (2022).

14. Li, Z., et al. Latent Diffusion Model for DNA Sequence Generation. arXiv [cs.LG] (2023).

15. Avdeyev, P., Shi, C., Tan, Y., Dudnyk, K. & Zhou, J. Dirichlet Diffusion Score Model for Biological Sequence Generation. ArXiv (2023) doi:10.48550/ARXIV.2205.15019.

16. Nichol, A. & Dhariwal, P. Improved Denoising Diffusion Probabilistic Models. (2021).

17. Ronneberger, O., Fischer, P. & Brox, T. U-Net: Convolutional Networks for Biomedical Image Segmentation. (2015).

18. Meuleman, W. et al. Index and biological spectrum of human DNase I hypersensitive sites. Nature 584, 244–251 (2020).

19. Meuleman, W. Synthetic DNA sequences. meuleman.org https://www.meuleman.org/research/synthseqs/.

20. Kent, W. J. BLAT--the BLAST-like alignment tool. Genome Res. 12, 656–664 (2002).

21. Korhonen, J., Martinmäki, P., Pizzi, C., Rastas, P. & Ukkonen, E. MOODS: fast search for position weight matrix matches in DNA sequences. Bioinformatics 25, 3181–3182 (2009).

22. Castro-Mondragon, J. A. et al. JASPAR 2022: the 9th release of the open-access database of transcription factor binding profiles. Nucleic Acids Res. 50, D165–D173 (2022).

23. Hansen, J. L. & Cohen, B. A. A quantitative metric of pioneer activity reveals that HNF4A has stronger in vivo pioneer activity than FOXA1. Genome Biol. 23, 221 (2022).

24. Lee, C. S., Friedman, J. R., Fulmer, J. T. & Kaestner, K. H. The initiation of liver development is dependent on Foxa transcription factors. Nature 435, 944–947 (2005).

25. Han, G. C. et al. Genome-Wide Organization of GATA1 and TAL1 Determined at High Resolution. Mol. Cell. Biol. 36, 157–172 (2016).

26. Wu, Y. et al. Highly efficient therapeutic gene editing of human hematopoietic stem cells. Nat. Med. 25, 776–783 (2019).

27. Yu, S. et al. Comprehensive analysis of the GATA transcription factor gene family in breast carcinoma using gene microarrays, online databases and integrated bioinformatics. Sci. Rep. 9, 1–16 (2019).

28. Reilly, S. K. et al. Direct characterization of cis-regulatory elements and functional dissection of complex genetic associations using HCR-FlowFISH. Nat. Genet. 53, 1166–1176 (2021).

29. Gasperini, M. et al. A genome-wide framework for mapping gene regulation via cellular genetic screens. Cell 176, 1516 (2019).

30. Argemi, J. et al. Defective HNF4alpha-dependent gene expression as a driver of hepatocellular failure in alcoholic hepatitis. Nat. Commun. 10, 1–19 (2019).

31. Miller, B. C. & Maus, M. V. CD19-Targeted CAR T Cells: A New Tool in the Fight against B Cell Malignancies. Oncol Res Treat 38, 683–690 (2015).

32. Pampari, A., et al. Bias factorized, base-resolution deep learning models of chromatin accessibility reveal cis-regulatory sequence syntax, transcription factor footprints and regulatory variants. (Zenodo, 2023). doi:10.5281/ZENODO.7567627.

33. Agarwal, V., et al. Massively parallel characterization of transcriptional regulatory elements in three diverse human cell types. bioRxiv (2023) doi:10.1101/2023.03.05.531189.

34. Avsec, Ž., et al. Effective gene expression prediction from sequence by integrating long-range interactions. Nat. Methods 18, 1196–1203 (2021).

35. Karollus, A., Mauermeier, T. & Gagneur, J. Current sequence-based models capture gene expression determinants in promoters but mostly ignore distal enhancers. Genome Biol. 24, 56 (2023).

36. Generating ‘smarter’ biotechnology. Nat. Biotechnol. 41, 157 (2023).

37. Ramesh, A. et al. Zero-Shot Text-to-Image Generation. in Proceedings of the 38th International Conference on Machine Learning (eds. Meila, M. & Zhang, T.) vol. 139 8821–8831 (PMLR, 18--24 Jul 2021).

38. Brown, T. B. et al. Language Models are Few-Shot Learners. arXiv [cs.CL] (2020).

39. Alamdari, S., et al. Protein generation with evolutionary diffusion: sequence is all you need. bioRxiv 2023.09.11.556673 (2023) doi:10.1101/2023.09.11.556673.

40. Watson, J. L. et al. Broadly applicable and accurate protein design by integrating structure prediction networks and diffusion generative models. *bioRxiv* 2022.12.09.519842 (2022) doi:10.1101/2022.12.09.519842.

41. The Annotated Diffusion Model. https://huggingface.co/blog/annotated-diffusion.

42. He, K., Zhang, X., Ren, S. & Sun, J. Deep Residual Learning for Image Recognition. (2015).

43. Shen, Z., Zhang, M., Zhao, H., Yi, S. & Li, H. Efficient Attention: Attention with Linear Complexities. (2018).

44. Ho, J. & Salimans, T. Classifier-Free Diffusion Guidance. NeurIPS 2021 Workshop on Deep Generative Models and Applications (2021).

45. Loshchilov, I. & Hutter, F. Decoupled Weight Decay Regularization. arXiv [cs.LG] (2017).

46. Avsec, Ž., et al. Base-resolution models of transcription-factor binding reveal soft motif syntax. Nat. Genet. 53, 354–366 (2021).

47. Quinlan, A. R. & Hall, I. M. BEDTools: a flexible suite of utilities for comparing genomic features. Bioinformatics 26, 841–842 (2010).

